# E3 ubiquitin-ligase Hakai induces LRP4 degradation and regulates Wnt/β-catenin signalling in colorectal cancer cells

**DOI:** 10.1101/2025.11.20.689197

**Authors:** Andrea Rodríguez-Alonso, Lía Jove, Gloria Alfonsín, Marta Tasende, Ingrid Jordens, Madelon Maurice, Angélica Figueroa

**Affiliations:** Epithelial Plasticity and Metastasis Group, Instituto de Investigación Biomédica de A Coruña (INIBIC), Complexo Hospitalario Universitario de A Coruña (CHUAC), Sergas, Universidade da Coruña (UDC), A Coruña, Spain; General Surgery and Digestive System Unit. Complexo Hospitalario Universitario de A Coruña (CHUAC), Sergas, A Coruña, Spain; Oncode Institute and Centre for Molecular Medicine, University Medical Centre Utrecht, Utrecht, the Netherlands

**Keywords:** E3 ubiquitin-ligase Hakai, LRP4, Wnt/β-catenin pathway, cancer stem cell, colon cancer

## Abstract

The epithelial-mesenchymal transition (EMT) is closely linked to the acquisition of cancer stem cell (CSC) properties, which contribute to treatment resistance and metastasis. This study investigates the role of the E3 ubiquitin-ligase Hakai, the first identified post-translational regulator of E-cadherin stability, in promoting CSC traits in colorectal cancer (CRC). To examine Hakai’s involvement in CSC regulation, we used an inducible shRNA in a HT29 cells. Under conditions that promote CSC characteristics, we silenced Hakai and evaluated tumoursphere formation and CSC marker expression. Proteomic and bioinformatic analyses were performed to identify Hakai-regulated proteins in tumoursphere cultures. Additionally, Western blot, RT-qPCR, co-immunoprecipitation, immunofluorescence and TOPFlash assays were employed to study CSC-related protein regulation in response to Hakai expression. Furthermore, we assessed the impact of Hakin-1, the pharmacological inhibitor specifically targeting Hakai’s HYB domain responsible for its E3 ubiquitin-ligase activity, on tumoursphere properties. Hakai silencing significantly reduced tumoursphere size and number accompanied by decreased expression of CSC markers and Wnt target genes. CSC-related proteins regulated by Hakai were identified, including LRP4, a negative regulator of Wnt/β-catenin pathway. Hakai interacts with LRP4, promoting its ubiquitination and degradation. Moreover, Hakai overexpression induces hyperactivation of Wnt/β-catenin sand disrupts LRP4’s inhibitory effect. Treatment with Hakin-1 effectively inhibited self-renewal and promoted differentiation within tumourspheres. These findings suggest that Hakai promotes CSCs properties by hyperactivation of the Wnt/β-catenin pathway via LRP4-mediated modulation. Additionally, Hakin-1 emerges as a promising therapeutic agent targeting CSCs by enhancing differentiation and attenuating Wnt/β-catenin activity, highlighting Hakai as a potential target for improving CSC treatment.

## Background

**Colorectal cancer (CRC)** is one of the most prevalent and lethal malignancies worldwide, characterized by high heterogeneity and frequent resistance to therapy. A growing body of evidence indicates that cancer stem cells (CSCs), a subpopulation of tumour cells with the ability to self-renew, differentiate, and tumour-initiating potential, play a critical role in tumour progression, metastasis, and resistance to therapy [1,2] Therefore, understanding the molecular mechanisms that regulate CSC properties is essential for developing more effective therapeutic strategies against CRC. Wnt/β-catenin signalling plays a critical role during homeostasis and in various diseases, including cancer. Indeed, over 90% of colorectal cancers involve hyperactivation of Wnt/β-catenin signalling [3,4]. The Wnt canonical pathway is dependent on β-catenin and TCF/LEF transcription factors, and primarily controls cell differentiation and proliferation. In the absence of Wnt ligands, the free pool of β-catenin is phosphorylated and incorporated into the β-catenin destruction complex (composed of APC, AXIN1/2, CK1, and GSK3β), leading to its degradation by proteasome. In the presence of Wnt ligands, the Wnt pathway is activated through Wnt binding to Frizzled (FZD) and LRP5/6 receptors. This binding leads to the activation of the cytosolic effector protein DVL, leading to inhibition of the destruction complex and thereby prevents β-catenin degradation. Subsequently, β-catenin accumulates in the cytoplasm and the excess translocates to the nucleus. Here, β-catenin activates TCF/LEF transcription factors, promoting the expression of Wnt/β-catenin target genes including *C-MYC*, *CCND1, AXIN2, MMP7* or *LGR5* [5,6]. The activation of Wnt/β-catenin target genes plays a crucial role in the initiation and maintenance of colorectal tumours and cancer stem cells (CSCs), making the Wnt/β-catenin pathway an attractive therapeutic target for CSC eradication [7]. On the other hand, LRP4 (also known as MEGF7), a member of the low-density lipoprotein receptor family, acts predominantly as a negative regulator of the Wnt/β-catenin pathway, often antagonizing LRP6-mediated signalling [8]. LRP4 function has been implicated in various neurological and developmental disorders. It plays a critical role in embryonic development, with loss-of-function mutations associated to malformation of limb, teeth, and kidneys [9–11]. It modulates both Wnt and BMP pathways, notably through interactions with antagonists like WISE, DKK1, and SOST [12,13]. Beyond Wnt, LRP4 influences other pathways, such as the PI3K/AKT pathway in epithelial–mesenchymal transition (EMT) [14]. Importantly, while emerging data hint at its role in cancer biology, LRP4’s precise contribution to Wnt/β-catenin signalling in colorectal cancer and its specific effects on CSC are poorly understood.

Ubiquitination is the second most prevalent post-translational modification after phosphorylation. This process involves the ATP-dependent covalent attachment of ubiquitin, a conserved protein composed of 76 amino acids, to a substrate protein through a cascade of enzymatic reactions involving the E1 activating enzymes, E2 conjugating enzymes, and E3 ubiquitin-ligase enzymes. E3 ubiquitin-ligases are pivotal regulators of protein stability and function, orchestrating the final step of the ubiquitination cascade that targets specific substrates for proteasomal degradation or modulates their activity, localization, and interactions [15,16]. Hakai was initially identified as an E3 ubiquitin-ligase that targets E-cadherin for ubiquitination, leading to its endocytosis and lysosomal degradation, a process that disrupts adherens junctions, facilitates cell–cell dissociation, and promotes EMT [17]. This classic function positioned Hakai as a critical regulator of epithelial plasticity [18]. Subsequent studies by our group and others have expanded its oncogenic profile, linking Hakai to enhanced tumour cell invasion, migration, and cellular plasticity across various cancer types [19–21]. Structurally, Hakai contains a RING finger domain typical of E3 ligases and a conserved Hakai HYB (Hakai pTyr-binding) domain, which mediates the recognition of phosphorylated substrates such as E-cadherin. Notably, the development of Hakin-1, a selective small-molecule inhibitor of Hakai, has opened new avenues for therapeutic intervention by targeting its E3 ligase activity [22].

The Wnt/β-catenin pathway is one of the key activators of the EMT, and it has been shown to play a role in the formation and maintenance of CSCs across various carcinomas [18,23]. It plays a pivotal role in the development and regeneration of the small intestinal epithelium, including the differentiation of Paneth cells at the crypt base [24]. In the intestinal and colonic epithelium, activation of Wnt target genes promotes the expansion of the stem cell compartment, whereas attenuation of Wnt/β-catenin signalling facilitates cellular differentiation [25] A functional relationship has been established between EMT and the acquisition of CSC properties, which is closely linked to therapy resistance, metastasis, and tumour relapse. Despite these advances, the role of Hakai in regulating CSC properties and its potential functional interplay with Wnt/β-catenin pathway remain largely unexplored.

In the present study, by using a colon cancer tumoursphere culture grown under conditions that promote the induction of CSC-like properties, we demonstrate that silencing Hakai reduces both the size and number of tumourspheres, accompanied by a decrease in CSC markers. Through proteomic and computational analyses, we identify several CSC-related proteins, notably LRP4. We show that Hakai interacts with LRP4, promoting its ubiquitination and subsequent degradation. Furthermore, Hakai overexpression activates Wnt/β-catenin signalling and cooperates with LRP6 to overcome the antagonistic effect of LRP4. Together, our findings uncover a novel regulatory mechanism by which Hakai modulates Wnt signalling via LRP4 degradation, with potential implications for CSC maintenance and colon cancer progression. Moreover, treatment with Hakin-1, a pharmacological inhibitor of Hakai, impairs tumoursphere formation by reducing stemness-associated features, disrupting 3D structural integrity, and promoting cellular differentiation, further supporting the therapeutic potential of targeting Hakai in colorectal cancer.

## Methods

### Reagents and antibodies

The primary antibody used for Western blot and immunofluorescence assays were as follows: Hakai (Invitrogen, 36-2800), LRP4 (Abcam, Ab174637), GAPDH (Invitrogen, 39-8600), LGR5 (Abcam, ab75850 and Invitrogen, MA5-25644), NANOG (Santa Cruz Biotechnology, sc-374001), Tubulin (Sigma-Aldrich, T9026), Vinculin (Cell Signalling Technology, E1E9V), MUC2 (Cell Signalling Technology, CCP58), LC3 I/II (Cell Signalling Technology, 4108), LRP6 (Cell Signalling, C47E12), V5 (Genscript, AØ1724 and Sigma, V8137), HA (Roche, 12CA5), E-cadherin (BD Bioscience, 610182 and Abcam, ab11512) and PRPS2 (Abcam, Ab96222). As secondary antibodies anti-mouse IgG (GE healthcare, NA934), anti-rabbit IgG (GE healthcare, NA931), anti-mouse-IgG-488 (Life Technologies, A11001), anti-rat-IgG-488 (Life Technologies, A11006), anti-mouse-IgG-594 (Life Technologies, A11005), anti-rat-IgG-568 (Life Technologies, A11077). Proteasome inhibitor MG132 (Sigma-Aldrich) was added for 6 h using 10 µM and 30 µM. Lysosome degradation inhibitor Chloroquine (Sigma-Aldrich), was employed for 24 h at 50 µM and for 6 h at 100 µM. Autophagy inhibitor 3-Methyladenine (Sigma-Aldrich) was added for 24h at 5 mM and 10 mM. Compound Hakin-1 [4-(5-{[2-(4-nitrophenyl)-2-oxoethyl]thio}-1H-tetrazol-1-yl)benzoic acid] were obtained from ChemBridge Corporation.

### Cell culture

Human colon cancer cell lines HCT116 (ATCC®#CCL-247TM) were obtained from the ATCC and were grown in DMEM. HEK293/T were cultured in Dulbecco’s Modified Eagle’s Medium (DMEM) - High Glucose (Gibco). Human colon adenocarcinoma HT29 cell line was purchased from Sigma-Aldrich and cultured in Mc Coy’s Modified Medium (Gibco). HT29 cells with a doxycycline-inducible lentiviral system for sh*CBLL1* was previously established by our group [26]. The doxycycline-inducible lentiviral system for *CBLL1* (SMARTvector Inducible Lentiviral sh*CBLL1*) was obtained from Dharmacon (Horizon Perkin Elmer Group). Lentiviral particles encoding shRNA for *CBLL1* or a non-targeting control (sh*CONTROL*) were generated and propagated following established protocols for transduction into cancer cells. HT29 sh*CONTROL* and HT29 sh*CBLL1* cells were cultured in McCoy’s 5A Medium with doxycycline (1 μg/mL; Sigma-Aldrich) to induce shRNA expression. Following induction, the efficiency of Hakai (*CBLL1 gene*) knockdown was verified by Western blot analysis. All culture medium were supplemented with 1% penicillin-streptomycin (Gibco, 5000 U/ml) and 10% of Fetal Bovine Serum (FBS, Corning) for monolayer cultures. Cells were cultured at 37 °C in a humidified incubator with a CO_2_ concentration of 5%. All cell lines were authenticated, used from early passages and monthly tested for mycoplasma contamination.

### Tumourspheres formation assays

Cells were placed into ultra-low attachment 6-well plates (Costar 6-well Clear Flat Bottom Ultra-Low Attachment Multiple Well Plates, Corning) at a density of 10 x 10^4^ cells per well in a Stem Cell Medium (20 ng/ml human EGF, 10 ng/ml human b-hFGF, 1x B27 and 1x GlutaMax in DMEM/F-12 Media), and then incubated at 37 °C with 5% CO_2_ from 3 to 5 days. Flat bottom plates were used to avoid the formation of cell aggregates instead of tumourspheres structures. Tumourspheres formation was monitored and characterized by phase-contrast microscopy using an Eclipse-Ti microscope (Nikon). After the incubation period, images of tumourspheres formed in each well were captured. To analyze tumoursphere size, area of at least 15 tumourspheres per experiment was measured using the Freehand Tool in ImageJ software (National Institutes of Health, USA). To quantify the number of tumourspheres, self-renewal assays were conducted as follows: cells were seeded at a density of 500 cells/mL in 24-well low-attachment plates, as previously described. After 5 days of culture, tumourspheres were analyzed by microscopy, and the number of tumourspheres was counted. The size and number of tumourspheres were graphically represented using GraphPad software.

### Western blot analysis

Cellular proteins were extracted using lysis buffer containing 1% Triton X-100, 20 mM Tris-HCl (pH 7.5), and 150 mM NaCl, supplemented with protease inhibitors: 10 μg/ml aprotinin, 10 μg/ml leupeptin, and 1 mM phenylmethylsulfonyl fluoride (PMSF). To prevent the activity of deubiquitinating enzymes during ubiquitination and co-immunoprecipitation assays, 10 mM N-ethylmaleimide (Sigma-Aldrich) was added. Protein concentration was determined using the Pierce™ BCA Protein Assay Kit (Thermo Fisher Scientific), following the manufacturer’s instructions. Equal amounts of protein were resolved by SDS-polyacrylamide gel electrophoresis (SDS-PAGE) and transferred onto polyvinylidene difluoride (PVDF) membranes (Millipore). Membranes were blocked for 1 hour at RT under agitation in blocking buffer [5% non-fat dry milk (Sigma-Aldrich) in TBS-T (20 mM Tris base, 150 mM NaCl, 0.1% Tween-20, pH 7.4)]. After blocking, membranes were cropped and incubated with specific primary antibodies to the proteins of interest, followed by incubation with appropriate HRP-conjugated secondary antibodies. Signal detection was performed using the Luminata™ Crescendo Western HRP Substrate (Millipore) and visualized with the Amersham Imager 600 system (GE Healthcare).

### RNA extraction and quantitative real-time PCR

Total RNA was extracted using TriPure Isolation reagent (Roche) and converted to cDNA using the NZY First-Strand cDNA Synthesis Kit (NZYTech). qRT-PCR was performed using LightCycler® 480 SYBR Green I Master (Roche) on an LightCycler® 480 Instrument (Roche). Comparative CT method (ΔΔCT method) was performed to analyse qPCR data using *β-ACTIN* as a housekeeping control. Primer sequences are detailed in **Table S1.**

### Proteomic analysis

Equal amounts of each sample were reduced with 10 mM dithiothreitol (DTT) for 1 hour at 37 °C under agitation. Subsequently, alkylation with 50 mM iodoacetamide (IAA) was performed in the dark for 45 minutes at RT. Samples were digested with sequencing grade-modified trypsin (Promega) at 1:30 enzyme-to substrate ratio. After 17 hours of digestion at 37 °C under agitation, samples were acidified with 10% trifluoroacetic acid to ∼pH 3 to stop the reaction and then, samples were concentrated using a under speed-vacuum (SpeedVac, Thermo Fisher Scientific) and sonicated for 3 minutes. Subsequently, the digested peptides were desalted using in-house made stage tips (3M Empore SPE-C18 disk, 47 mm, Sigma-Aldrich).

The dried eluates were reconstituted in water containing 0.1% FA for direct liquid chromatography-mass spectrometry (LC-MS) analysis. 200 ng of peptide mixture were loaded in a nanoElute (Bruker Daltonics) nano-flow LC that was coupled to a high resolution timsTOF Pro (Bruker Daltonics) with a CaptiveSpray ion source (Bruker Daltonics). Liquid chromatography was conducted at 50 °C with a constant flow of 400 nL/minute on a reversed-phase column (15 cm x 75 m i.d.) with a pulled emitter tip, packed with 1.9 m C18-coated porous silica beads (Dr. Maisch, Ammerbuch-Entringen, Germany). Chromatographic separation was performed using a linear gradient of 5-35% Buffer B (100% acetonitrile and 0.1% FA) over a period of 30 minutes, followed by an increase to 95% within 2 minutes and maintaining at 95% for 8 minutes. To prevent the sample carryover, two blank injections were performed after each sample run to prevent any residual material. This involved a 30 minutes short gradient total run time (transitioning from 2% to 95% of Solvent B over 5 minutes; then from 95% to 5% B during another 5 minutes; maintaining 5% B for 5 minutes, reaching 95% at 25 min, and finally holding at 95% for 5 minutes to wash the column). Peptides underwent electrospray ionization (ESI) and were analyzed using data-dependent acquisition (DDA) mode with parallel accumulation–serial fragmentation (PASEF) enabled.

Raw data files were analyzed to obtain protein identifications using MaxQuant and LFQ Analyst was used for the analysis of Label-Free Quantitative (LFQ) proteomic data sets. Statistical analyses were carried out with an in-house R script using the “MaxQuant “proteinGroup.txt” as the primary input file in conjunction with an experimental design text file, which describes conditions and replicates. Contaminant proteins, reverse sequences, proteins identified “only by site” or with “only by a single peptide” were removed. LFQ intensities data were converted to a log2 scale, and replicates were grouped by conditions based on the information provided in the “experimental design.txt” file. Missing values were imputed using the “Missing not At Random” (MNAR) method, which uses random draws from a Gaussian distribution left-shifted by 1.8 SD (standard deviation) with a width of 0.3. To determine significantly regulated proteins between conditions, a cutoff of an adjusted p-value cutoff of 0.05 (Benjamini−Hochberg method) along with a fold change of 1.5 and a requirement 74 of at least two peptides was applied.

### Plasmid transfection

Cell lines were transfected with plasmids pcDNA3.1, pcDNA-FLAG-Hakai and pBSSR-HA-Ubiquitin and which were kindly provided by Yasuyuki Fujita (Hokkaido University, Japan). pcDNA-LRP4 were a kindly gift of Tatsuo Suzuki (Shinshu University School of Medicine, Japan). Plasmids for TOPFlash, FOPFlash, TK-Renilla, Myc-LRP6, MESD and Myc-LGR5 were kindly provided by Madelon Maurice (Utrecht, Netherlands). The transfection experiments were performed using FuGENE® 6 Transfection Reagent (Promega) and Opti-MEM™ Reduced Serum Medium (Thermo Fisher), following the manufacturer’s protocol.

### Immunoprecipitation and ubiquitination assays

For immunoprecipitation experiments, cells were lysed using lysis buffer containing 1% Triton X-100, 20 mM Tris-HCl (pH 7.5), and 150 mM NaCl, supplemented with 10 μg/ml leupeptin, 10 μg/ml aprotinin, 1 mM PMSF and 10 mM N-ethylmaleimide for 30 minutes at 4 °C. After centrifugation at 18,000×g for 10 minutes, supernatants were immunoprecipitated with 2.5 μg of anti-LRP4 antibody or mouse IgG (Santa Cruz Biotechnology, USA) at 4 °C for 2 hours, bound to 60 µL protein G PLUS-Agarose beads (Santa Cruz Biotechnology). Immunoprecipitated proteins and input control protein lysates were then loaded into SDS-PAGE gels to perform Western blot analysis with the indicated antibodies.

### TOPFlash luciferase reporter assay

HEK293T or HCT116 cells were seeded into 24-well plates and grown overnight. The next day, cells were transfected with 30 ng of the reporter plasmid TOPFlash or FOPFlash, 5 ng Thymidine Kinase (TK)-Renilla and 50-150 ng of each of the indicated plasmids. Plasmids concentrations were 150 ng pcDNA-V5-Hakai, 50 ng pcDNA-LRP4, 100 ng MYC-LRP6 + 25 ng MESD and 100 ng MYC-LGR5, respectively. All transfections were performed with FuGENE® 6 Transfection Reagent according to the manufacturer’s protocol. Six hours post transfection, cells were incubated with control L-cell conditioned medium (LCM) or WNT3A-conditioned medium (WCM) overnight. WNT3a-conditioned medium (CM) were produced as described [27]. After 24 hours, total cell lysates were extracted with Passive Lysis Buffer (Promega) for 20 minutes at room temperature and levels of Renilla and Firefly luciferase was measured using the Dual-Luciferase kit (Promega) according to manufacturer’s instructions using a Centro LB960 luminometer (Berthold).

### Immunofluorescence assays

For immunofluorescence experiments of cells, HEK293T cells were seeded on laminin-coated glass coverslips in 24-well plates at a density of 0.25 cm^2.^ After overnight incubation, cells were transfected with 150 ng PCDNA-V5-Hakai, 100 ng PCDNA-LRP4 and/or empty vector PCDNA3.1 during 24 hours. Transfection was carried out using FuGENE® 6 Transfection Reagent and with a DNA ratio of 5:1. The next day, cells were fixed in 4% PFA for 30 minutes at room temperature. After the incubation, the reaction was quenched in 0.05 M NH_4_Cl for 15 minutes. Cells were blocked in PBS containing 2% BSA and 0.1% Saponin. Primary and secondary antibody incubations were performed in blocking buffer for 1 hour and 45 minutes at room temperature, respectively. Cells were mounted in ProLong Gold (Life Technologies) and left drying overnight in the dark before being analysed using an LSM700 confocal microscope.

To perform immunofluorescence assays of tumourspheres, tumourspheres culture were collected and centrifuged. After washing with cold PBS, tumourspheres of each ultra-low attachment 6-well were seeded in one well of an 8-well chamber (Millicell EZ SLIDE 8-well glass, Millipore) and they were fixed in 200 μl of 4% PFA in PBS for 40 minutes at room temperature, then permeabilized and blocked with PBD 0.2 T buffer [1% BSA, 1% DMSO, 0.2% Triton X-100 in PBS] for 1 hour. Primary antibodies were incubated overnight at 4°C, followed by incubation with secondary antibodies for 3 hours in the dark. Nuclei were stained with Hoechst 33342 (Life Technologies) 1:5000 diluted in PBS for 5 minutes and tumourspheres were mounted for microscopy using ProLong Gold antifade reagent (Life Technologies). Images were acquired with a Nikon A1R confocal microscope and analysed using Python with NumPy and Matplotlib to measure fluorophore intensities. Results were quantified from five pictures in three replicate experiments, expressed as mean ± SEM, and analysed by t-test for statistical significance.

### Statistical analysis

Results are presented as mean ± SEM, as indicated in the figures. Statistical analysis and graphical representations were performed using GraphPad Prism (Version 8, GraphPad Software). The Shapiro-Wilk test was applied to assess the normality of the data and determine whether they followed a Gaussian distribution. Statistical significance was evaluated using a t-test for comparisons between two groups, or ANOVA for comparisons involving three or more groups. If the ANOVA revealed significant differences, post-hoc tests were conducted to identify which groups differed significantly. Dunnett’s post-hoc test was used when comparing multiple treatment groups to a single control. The results are presented in the figures as *p < 0.05, **p < 0.01, ***p < 0.001, and ****p < 0.0001.

## Results

### Hakai-silencing reduces colon cancer tumourspheres and modulates self-renewal and differentiation-related properties

To investigate the mechanistic insights of Hakai in modulating colon CSCs, we used HT29 tumourspheres, a three-dimensional culture model derived from the human colon adenocarcinoma cell line HT29. This model has been extensively used to study CSCs within the context of colorectal cancer [28]. HT29 tumourspheres serve as a valuable tool for advancing our understanding of CSC biology and developing novel therapeutic approaches for targeting CSCs in colorectal cancer and other tumour types. Hakai-silencing was induced in an HT29 monolayer culture by using a previously reported inducible viral-transduced system by doxycycline [26]. After 72 hours, HT29 tumourspheres were induced for 5 days in ultra-low attachment plates with a specific medium to enrich CSC growth, as shown in the schematic workflow (**Fig. 1A**). This assay showed that Hakai-silencing reduces the number and size of the tumourspheres (**Fig. 1B-D**), further indicating the impact of Hakai on cancer cells self-renewal capacity and proliferation. In accordance with previous reports [29], Hakai-silencing was accompanied by the downregulation of stem cell markers and Wnt/β-catenin targets genes (**Fig. 1E-F**) including LGR5, the best-established CSC biomarker for colorectal, and NANOG transcription factor, a universal CSC marker. These findings highlight the contribution of Hakai to the development of CSC tumourspheres, reinforcing its potential relevance as a therapeutic target in colorectal cancer.

**Fig 1.**
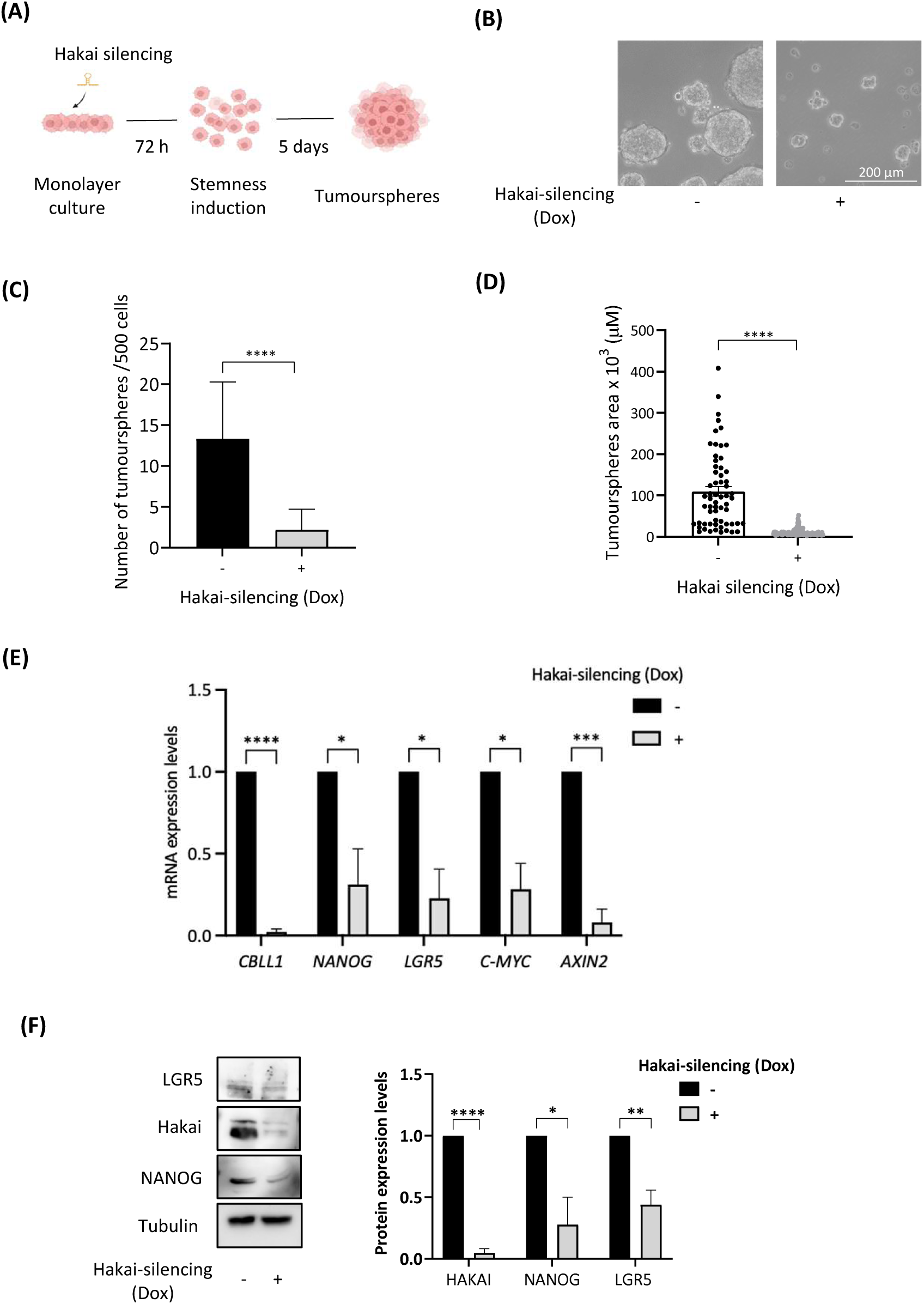
Hakai-silencing results in a decrease in the number and size of tumourspheres while decreasing stem cell markers. **(A)** Schematic workflow of the self-renewal model. Hakai-silencing was induced in HT29 monolayer cultures and after 72 hours, cells were cultured in cancer stem cell promoting conditions for 5 days to form tumourspheres. **(B)** Tumourspheres were phenotypical characterized by phase contrast images. Representative images of Hakai-silenced HT29 tumourspheres and control conditions were taken after 5 days of the induction of stemness. Images were obtained using a 10x objective. Scale bar, 200 µm. **(C)** Quantification of the number of HT29 tumourspheres in Hakai-silenced compared to control conditions. (D) Size of tumourspheres of HT29 tumourspheres in Hakai-silenced compared to control conditions. Results are expressed as mean ± SEM. Quantification was carried out using ImageJ software, and statistical significance was determined using GraphPad Prism software. **(E)** mRNA expression levels of Hakai (CBLL1 gene) and stem cell markers in HT29 tumourspheres in Hakai-silenced compared to control analysed by RT-PCR. **(F)** Expression of Hakai and stem cell protein markers was determined by Western blot in HT29 tumourspheres with Hakai silencing compared to control. Protein bands were quantified using ImageJ and normalized to loading control. Data are presented as mean ± SEM from three independent experiments. Statistical significance was determined using an unpaired t-test (*p < 0.05, **p < 0.01, ***p < 0.001, ****p < 0.0001).

### Proteomic and bioinformatic analysis identifies novel cancer-related candidates regulated by Hakai

In order to investigate the extent to which Hakai-silencing may control altered protein expression in CSC, we decided to follow two different strategies: a proteomic study comparing Hakai-silencing in colon cancer tumourspheres to control conditions, and consulting the UbiBrowser database that is able to predict the proteome-wide human Hakai-substrate interaction network [30]. For the proteomic analysis we used nano-LC-MS/MS analysis coupled to a timsTOF Pro mass spectrometer to identify novel Hakai-regulated proteins in colon CSC tumourspheres. The schematic workflow of the process is represented (**Fig. 2A**). The confirmation of Hakai-silencing was shown at the protein level in three biological replicates per experimental group analysed in the proteomic study (**Fig. 2B**). Mass spectrometry analysis identified a total of 3,602 proteins with ≥ 2 peptides. A total of 103 proteins (2.85%) were specifically enriched in Hakai-silenced HT29 tumourspheres (+ Dox), whereas 118 proteins (3.28%) were specifically enriched in control HT29 tumourspheres (-Dox) (**Fig. 2C**). The principal component analysis (PCA) and Volcano plot of differentially proteins identified are shown (**Fig. 2D-E**). Notably, the sum of the principal component PC1 (PC1, X-axis) and PC2 (PC2, Y-axis) captured the majority of the variance, approximately 80%, providing a good representation of the variance in the samples (**Fig. 2D**). The distinct clustering observed suggests significant differences in the proteome between the two conditions. We conducted a hierarchical clustering analysis of the differentially protein expression from three replicates comparing Hakai-silenced to control tumourspheres, which indicates protein expression was similar in each group (**Fig. 2E**). As represented by black dots in the Volcano plot analysis (**Fig. 2F**), eight proteins were significantly upregulated (Fold change ≥ 1.5 and p-value < 0.05) in Hakai-silenced tumourspheres, while three proteins were significantly downregulated. The boxplot represents the regulated proteins in Hakai-silenced tumourspheres compared to control tumourspheres (**Fig. 2G**), including MUC2, DDX55, CLMN, C19ORF43, RABGGTA, SLC35F2, UBXN7, and UFSP2 that were significantly enriched in Hakai-silenced tumourspheres, whereas NIT1, PRPS2, and WDR26 were significantly depleted. The complete dataset from the proteomic analysis is shown in **Table 1**, including the biological functions of the identified proteins obtained from Uniprot. Interestingly, several identified proteins are cancer-related proteins involved in CSC or differentiation. However, due to the characteristics of the ubiquitination cascade, which involves transient interactions characterized by rapid kinetics and weak target affinities, in general, E3 ubiquitin-ligase substrates were not identified, making it challenging to identify and study their specific substrates. Therefore, we used the UbiBrowser platform, which allowed us to determine the predicted Hakai-substrate interaction network [30]. By using UbiBrowser 2.0 [31], 116 proteins were identified as potential predicted substrates for Hakai-mediated ubiquitination (**Table S2)** [32]. We shortlisted the top 20 predicted novel substrates that could potentially be ubiquitinated by Hakai protein (in blue). Moreover, 4 already known substrates were identified, shown in red (**Fig. 2H**). Thus far, very few substrates for Hakai were biochemically or functionally reported, including E-cadherin, Cortactin, DOK1 or Ajuba and specific substrates of Hakai in CSC conditions remain unknown [17,20,33].

**Fig 2.**
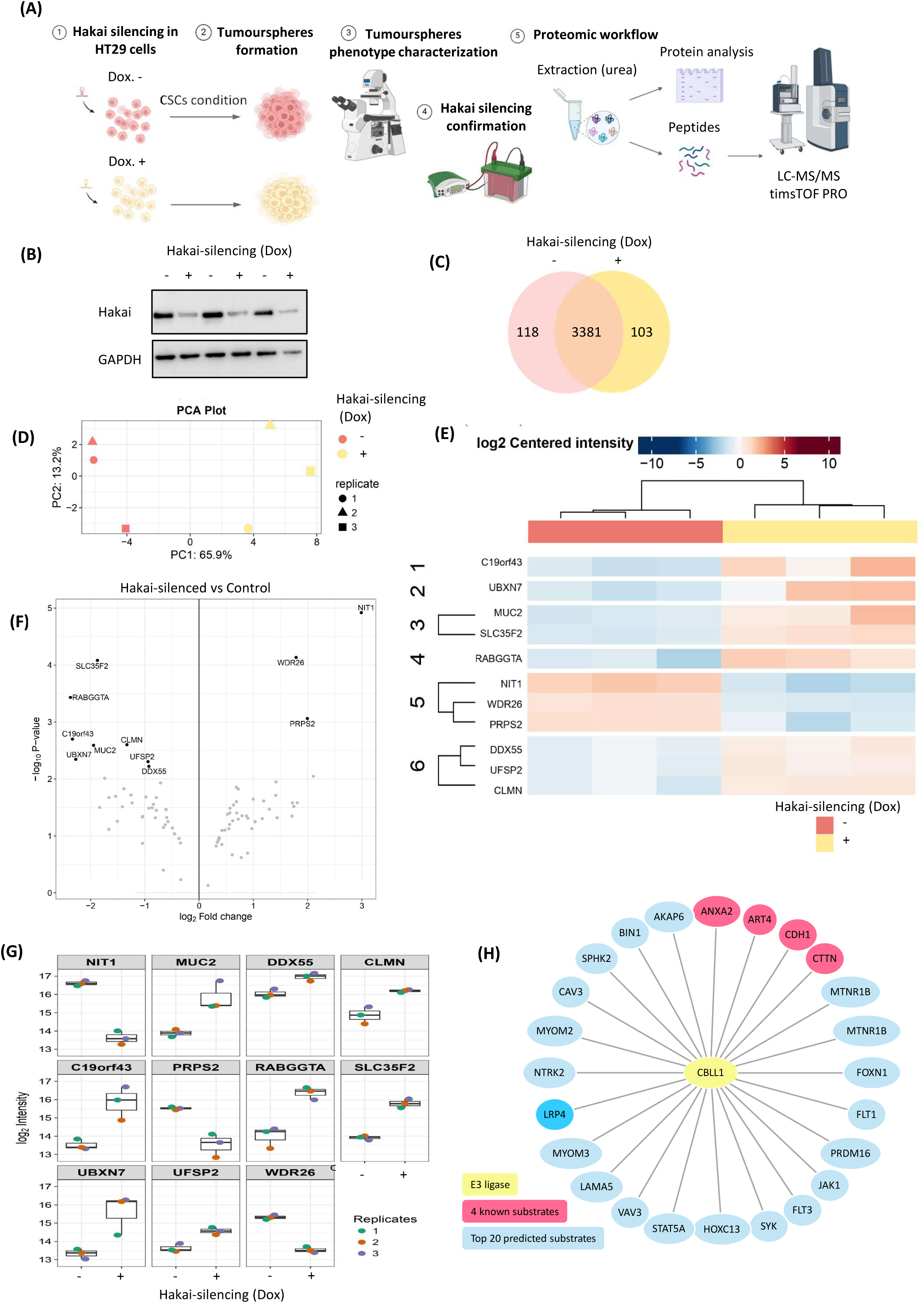
Proteomic study and UbiBrowser database to find novel Hakai-regulated proteins in tumoursphere formation. **(A)** Schematic workflow of the proteomic study in Hakai-silenced tumoursphere compared to control. Hakai-silencing in HT29 cells was induced and cells were seeded in stem cell promoting media using ultra-low attachment plates to induce tumoursphere formation. Protein extracts were digested and protein profile was analysed by silver staining in SDS-page gels. Proteins were digested with trypsin and the obtained peptides were fractionated and separated by nano-LC-MS/MS analysis coupled to a timsTOF Pro mass spectrometer. **(B)** Hakai-silencing in tumourspheres was confirmed in three biological replicates by Western blot. GAPDH was used as loading control. **(C)** Venn diagram of identified proteins (≥ 2 peptides) in tumourspheres obtained from Hakai-silenced HT29 colon cancer tumourspheres versus control. **(D)** Principle Component Analysis (PCA) plot showing two principal components corresponding to three replicates of Hakai-silenced tumourspheres and three replicates of control tumourspheres. **(E)** Heatmap showing the regulated proteins. Up and down regulated proteins in Hakai-silenced tumoursphere versus control are represented. **(F)** Volcano plot of differentially expressed proteins in Hakai-silenced tumourspheres versus control tumourspheres. Black dots represent proteins showing significant fold changes. The protein significance was set to adjusted p-value < 0.05, protein fold change to ≥ 1.5, used peptides to ≥ 2. **(G)** Box plot showing the protein profiling comparison, based on the label-free quantitation (LFQ) for which levels changed significantly between samples. The LFQ values are plotted on a Log2 scale along the vertical axis. **(H)** Top 20 Hakai (CBLL1 gene) substrates predicted in Homo sapiens (blue) and the 4 known substrates already described (red) by using UbiBrowser 2.0 database.

**Table 1.**
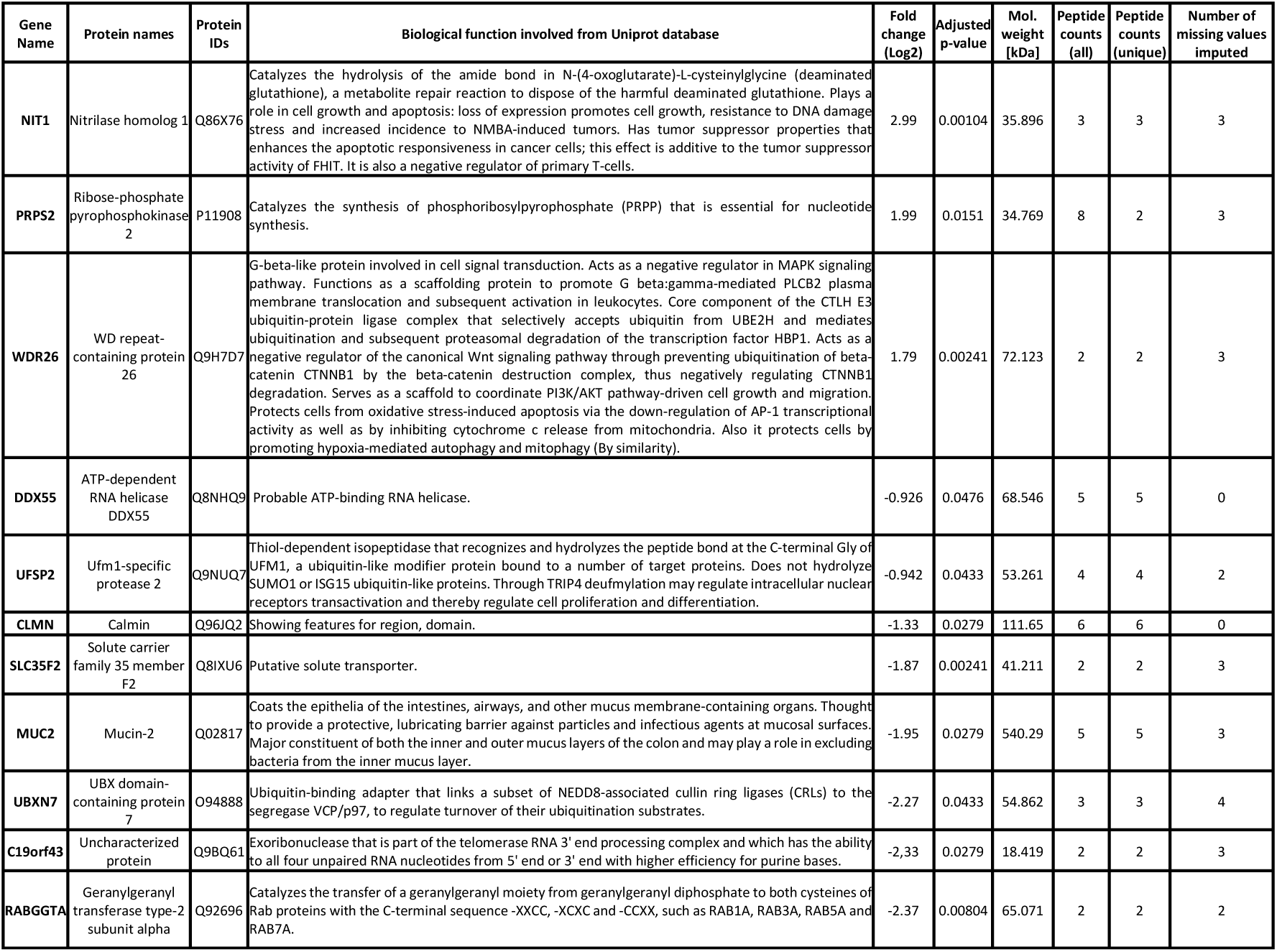
Identification of Hakai regulated proteins in Hakai-silenced HT29 colon cancer tumourspheres. Statistically significant identified proteins by iTRAQ analysis of Hakai-silenced HT29 colon cancer tumourspheres compared to control tumourspheres. The selected unique proteins found identified with fold change >1.5 and a cut-off of adjusted p-value < 0.05.

### Hakai interacts with LRP4 thereby inducing its ubiquitination and degradation

The predicted novel Hakai substrates identified by UbiBrowser included several cancer-related proteins. Given that previous studies have reported Hakai is involved in EMT and that our results support Hakai implication in CSC maintenance (**Fig. 1**), we directed our focus towards proteins involved in CSC regulation. Interestingly, our top four protein, LRP4 is a member of the low-density lipoprotein receptor (LRP) and can act as negative regulator of Wnt signalling [10]. LRP4 is primarily known in neuromuscular junction development and bone formation [10,12], however the significance in CSC is not well understood. To investigate the potential link between Hakai and LRP4, FLAG-Hakai was transiently transfected into HEK293 and colon cancer HCT116 cells, showing that Hakai overexpression reduced LRP4 protein levels in both cell lines (**Fig. 3A**). Next, immunofluorescence analyses were conducted using HEK293 cells transiently overexpressing V5-Hakai in presence or absence of overexpressed LRP4. Hakai overexpression alone was detected mainly in the nucleus but also in the cytosol, consistent with previous reports [34]. LRP4 was located at cell membrane, however, when both proteins were overexpressed, LRP4 expression was decreased (indicated by arrows) (**Fig. 3B**), indicating that Hakai may regulate LRP4 expression. As previously mentioned, the absence of co-localization of both proteins may be attributed to transient interactions between Hakai and LRP4. Given that Hakai overexpression downregulates LRP4 protein levels, and considering that LRP4 is a predicted new substrate for Hakai by UbiBrowser, we investigated whether we could detect a protein–protein interaction by immunoprecipitation. As co-immunoprecipitation using endogenous proteins was challenging, FLAG-Hakai, LRP4, HA-ubiquitin were transiently transfected in HEK293T cells and LRP4 antibody was used for immunoprecipitation. As expected, co-immunoprecipitation was observed for Hakai and LRP4 when overexpressed, further confirming Hakai and LRP4 interaction (**Fig. 3C**). As previous studies reported Hakai as an E3 ubiquitin-ligase that induces ubiquitination and subsequent degradation of its substrate, we predicted that LRP4 could act as a substrate for Hakai. To prove this hypothesis, we carried out an ubiquitination assay following the immunoprecipitation of LRP4. Western blot analysis showed that Hakai overexpression induces LRP4 ubiquitination, shown as a HA-Ub smear in LRP4 immunoprecipitates (**Fig. 3D**). Taken together, these results show that Hakai interacts with LRP4 and induces its ubiquitination. Then, we further investigated the mechanism by which LRP4 degradation was achieved by Hakai. We analysed the effect on LRP4 protein expression in presence or absence of the proteasome inhibitor MG132. We observed that LRP4 protein levels were increased in presence of MG132, without affecting the levels of Hakai (**Fig. 3E**). However, this effect was not observed in presence of lysosome inhibitor Chloroquine nor in the presence of the autophagy inhibitor 3-MA (**Fig. S1**). Notably, MG132 treatment in Hakai-overexpressing cells resulted in the restoration of LRP4 levels (**Fig. 3F**), further supporting that the induction of LRP4 degradation by Hakai occurs, at least partially, in a proteasome-dependent manner. Moreover, treatment of cells with the previously reported Hakai inhibitor, Hakin-1 that specifically blocks the HYB domain responsible for the E3 ubiquitin-ligase activity of Hakai, was also able to rescue downregulation of LRP4 induced by Hakai in HCT116 cells (**Fig. 3G**). Thus, our results support that Hakai mediates LRP4 degradation through its ubiquitin-ligase activity, further supporting LRP4 as a novel substrate for Hakai.

**Fig 3.**
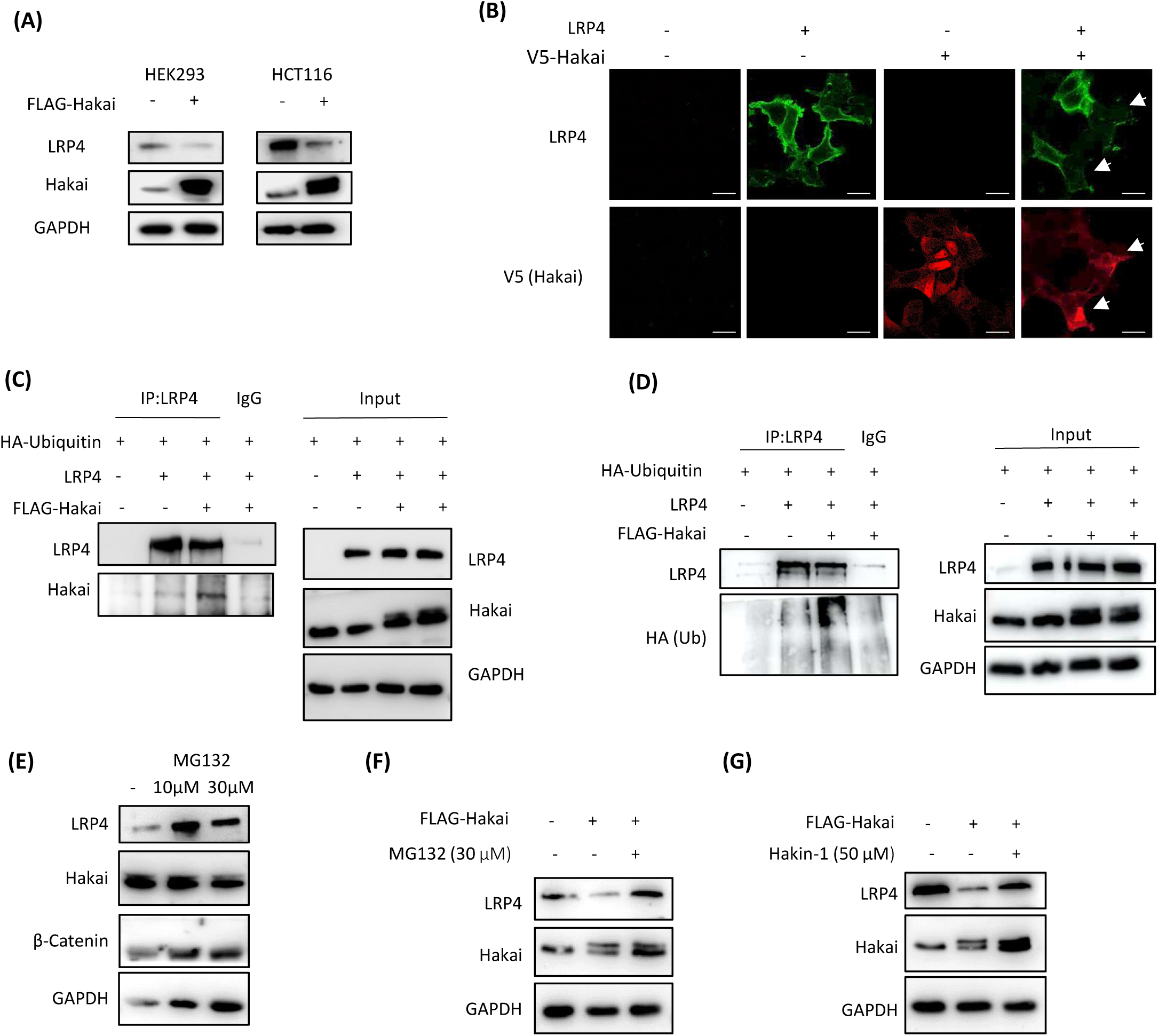
Identification of LRP4 as a new potential Hakai-interacting protein. **(A)** Hakai overexpression reduces LRP4 protein levels in HEK293 and HCT116 cells shown by Western blot. **(B)** Impact of Hakai on cell-surface expression of LRP4 in HEK293T transfected with LRP4 and empty vector or Hakai-V5. Surface levels of LRP4 were detected in unpermeabilized cells after which total levels of Hakai were visualized via permeabilization using confocal microscopy. Pictures were taken using a 63x objective and scale bar represents 30 µm. **(C)** Coimmunoprecipitation of Hakai and LRP4 in HEK293 cells overexpressing FLAG-Hakai, LRP4 and HA-ubiquitin. **(D)** Hakai-dependent ubiquitination of LRP4. FLAG-Hakai, HA-ubiquitin and LRP4 were transiently transfected into HEK293 cells. Immunoprecipitation was performed with anti-LRP4 and analyzed by Western blot. **(E)** LRP4 and FLAG-Hakai levels in HCT116 cells treated with the proteasome inhibitor MG132 analysed by Western blot. β-catenin were used as a positive control for MG132 treatment. **(F)** HCT116 cells were transiently transfected with FLAG-Hakai and the next day treated with MG132 (30 µM) for 6 h. Endogenous LRP4 levels were analysed by Western blot. **(G)** HCT116 cells were transiently transfected with FLAG-Hakai and treated with 50 µM of Hakin-1. LRP4 endogenous levels were analysed by Western Blot.

### Hakai counteracts the antagonistic effect of LRP4 on LRP6-mediated Wnt pathway activation

LRP4 is a transmembrane protein of the LDL receptor family and functions as a negative regulator of the canonical Wnt/β-catenin signalling pathway by antagonizing LRP6 receptor [8]. Considering the crucial role of LRP4 in development and its implication in the Wnt/β-catenin pathway, we aimed to investigate the potential mechanism by which Hakai may influence Wnt/β-catenin signalling. Moreover, the effect of Hakai on key Wnt-related proteins was extended to LRP6. First, we analysed the effect on β-catenin-TCF transcriptional activity by using the TOP-Flash reporter gene, in which the luciferase gene is placed under the control of a promoter harbouring ten copies of the TCF/LEF-1 consensus responsive sequence with a c-fos minimal promoter. As negative control, a mutated form of this promoter (FOP-Flash) was used. Cells were transfected with either TOP-Flash or FOP-Flash plasmids and the ratio of luciferase activity from TOP-Flash to FOP-Flash was calculated to obtain a measurement of Wnt-specific transcriptional activity. TK-Renilla was transfected as a transfection control. Cells were co-transfected with the Hakai and LRP4 or LRP6 plasmids (**Fig. 4A-B**). We added either control L-cell conditioned medium (LCM) or WNT3A-conditioned media (WCM) to study the effects of Hakai on both basal as well as Wnt-stimulated conditions. As shown, overexpression of LRP4 significantly inhibited both basal WNT signalling as well as WNT3A-induced signalling, in line with LRP4 negatively regulating Wnt/β-catenin signalling. Whereas in basal conditions overexpression of Hakai showed no significant effect, Hakai significantly potentiated WNT3A-induced signalling. However, Hakai was unable to fully rescue the inhibitory effect of LRP4 when both were co-transfected, although a small effect on basal LRP4-mediated suppression was observed (**Fig. 4A**). On the other hand, we confirmed that LRP6 was able to significantly activate Wnt-pathway in both basal and WNT3A-stimulated conditions [35]. Importantly, there was a synergistic effect observed when Hakai and LRP6 were co-overexpressed. LRP4 co-expression was able suppress LRP6-induced WNT pathway activation, suggesting that LRP4 acts as an antagonist of LRP6 (**Fig. 4B**). Interestingly, co-expression of LRP4, LRP6, and Hakai showed that Hakai was able to partially rescue/counteract the negative effect of LRP4 on LRP6, potentially by inducing the degradation of LRP4 (**Fig. 4B)**.

**Fig 4.**
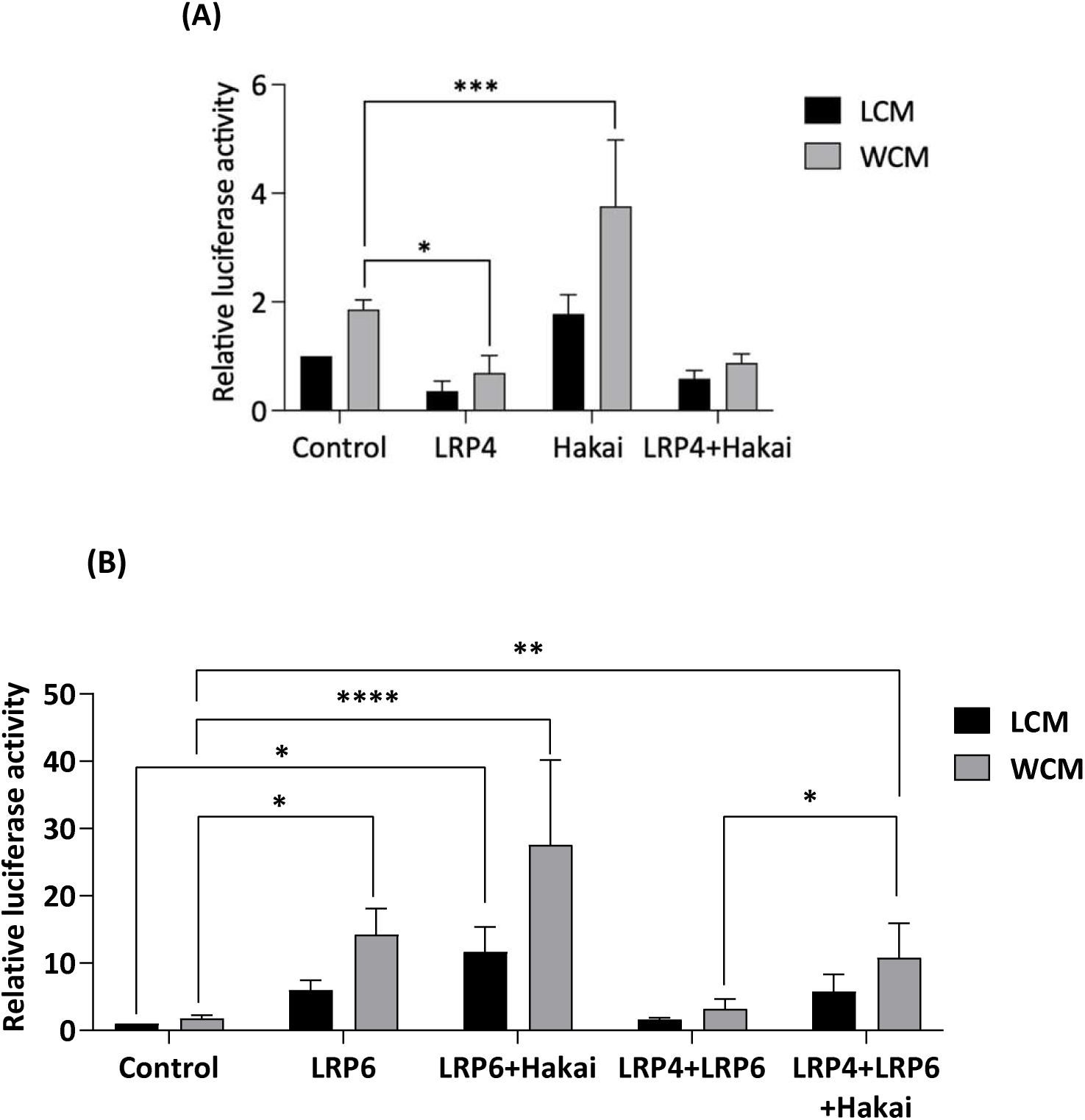
Hakai modulates β-catenin–TCF/LEF-1 signalling. Wnt signalling activity upon overexpression of the indicated plasmid in colon cancer HCT116 treated with control L-cell conditioned medium (LCM) and WNT3A conditioned medium (WCM). Cells were transfected with TOP-flash or FOP-flash reporter plasmids together with the indicated plasmids **(A)** Hakai-V5 and/or LRP4; **(B)** Hakai-V5 and/or LGR5 and **(C)** Hakai-V5, LRP4 and/or LRP6. Relative luciferase activity is presented as fold of control in mean ± SEM of three independent experiments.Two-ways ANOVA with Dunnet’s post hoc analysis was performed for statistical analysis (*p < 0.05, **p < 0.01, ***p < 0.001, ****p < 0.0001).

### Hakin-1 disrupts tumourspheres formation and stemness

Preclinical studies from our group have demonstrated the therapeutic potential of targeting Hakai through its HYB domain with Hakin-1 [22]. We therefore next investigated the potential effect of pharmacological inhibition of the E3 ubiquitin ligase Hakai using Hakin-1 on tumoursphere formation. HT29-derived tumourspheres were treated with Hakin-1 for six days, concurrently with the induction of stemness culture conditions (**Fig. 5A**). Phase-contrast microscopy revealed that this treatment phenotypically affected the tumourspheres (**Fig. 5B**). Treated tumourspheres exhibited a less compact morphology and poorly defined 3D structures compared to the well-organized and dense tumourspheres observed under control conditions, suggesting a potential loss of stemness and increased cellular differentiation. In addition, Hakin-1 modestly, but significantly, reduced the number of tumourspheres formed (**Fig. 5C**). Although the area of individual tumourspheres varied, Hakin-1 treatment significantly decreased the average size of tumourspheres compared to the control condition (**Fig. 5D**). Importantly, similar effects were observed when Hakin-1 treatment was applied either before or after the induction of stemness. To further investigate the effect of Hakin-1 on stemness and differentiation, we analysed the expression levels of various markers for the Wnt/β-catenin pathway, stem cells, and differentiation in tumourspheres. Hakin-1 significantly reduced the expression levels of *LEF-1* and *TCF-1*, both of which play pivotal roles in the Wnt/β-catenin signalling pathway through their interaction with β-catenin. Moreover, Hakin-1 markedly reduced the expression of the stem cell markers *LGR5* and *NANOG* **(Fig. 5E),** while there was no significant reduction in the expression of downstream Wnt/β-catenin target genes commonly upregulated in colorectal cancer (*CCND1*, *C-MYC*, and *MMP7*) (**Fig. 5F**). Taken together, Hakin-1 impairs Wnt pathway activation, particularly affecting genes related to stemness. At the protein level, we observed a slight upregulation of E-cadherin, accompanied by a downregulation of Wnt-related stem cell markers LGR5 and NANOG (**Fig. 5G**). As expected, Hakin-1 treatment did not alter Hakai protein levels. These findings support the notion that Hakai inhibition alters the tumourspheres phenotype by reducing stemness-associated markers.

**Fig 5.**
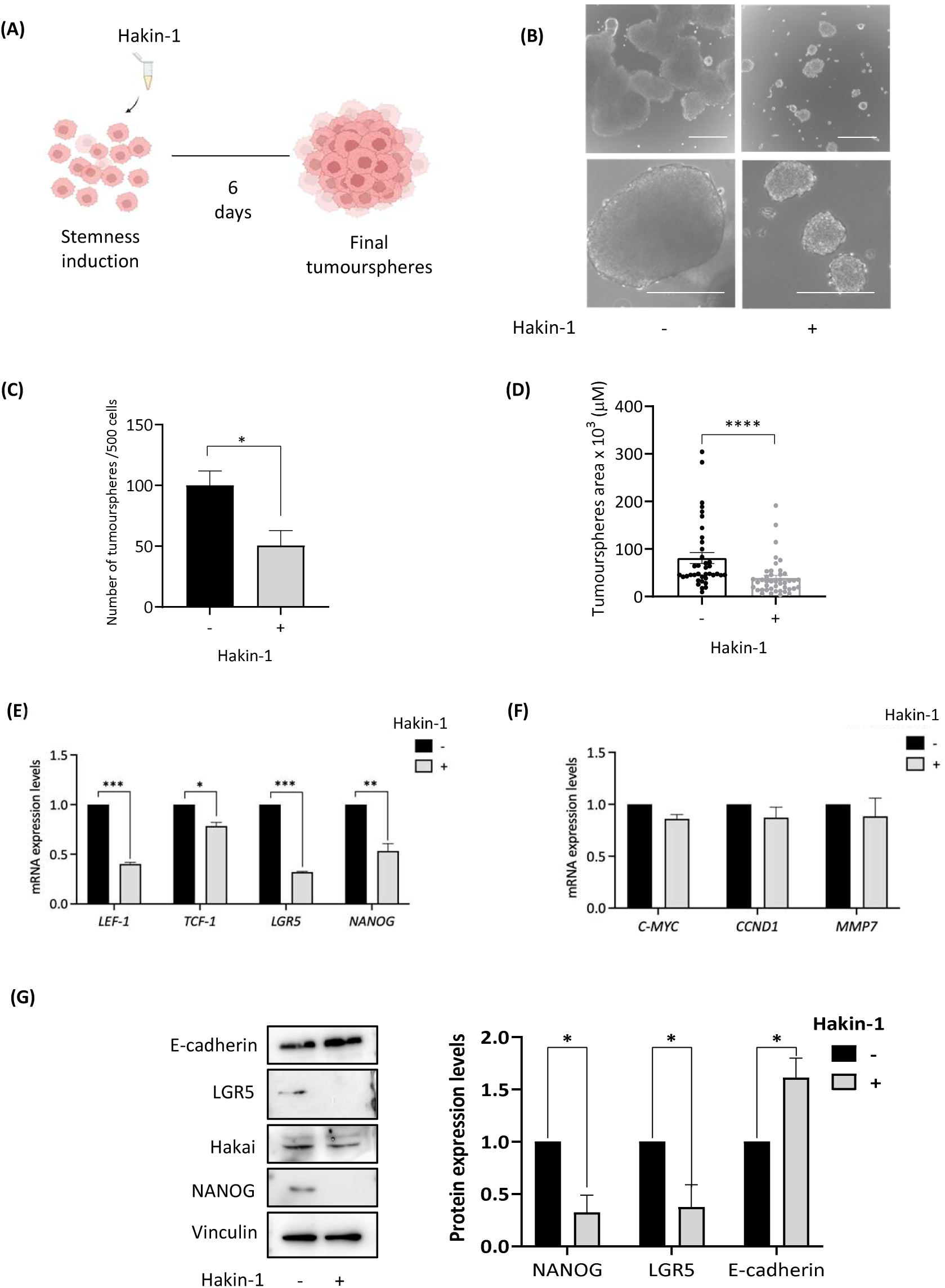
Impact of Hakin-1 on tumoursphere formation. **(A)** Schematic representation of the experimental workflow used to assess self-renewal. Hakin-1 (50 µM) was added at the initiation of cancer stem cell induction. **(B)** Representative images of HT29 tumourspheres treated with Hakin-1 or DMSO (vehicle control) five days after induction under stemness-promoting conditions. Images were acquired using a using a 4x (upper images, scale bar: 250 µm) or 10x (lower images, scale bar: 200 µm) objective. **(C)** Quantification of the number of tumourspheres formed from HT29 cells treated with Hakin-1 or DMSO. **(D)** Quantification of tumoursphere size in the same conditions. Data in (C) and (D) are presented as mean ± SEM. Quantification was performed using ImageJ software, and statistical analysis was conducted with GraphPad Prism. **(E-F)** Effect of Hakin-1 on Wnt/β-catenin signalling and stem cell markers at mRNA levels. RT-qPCR analysis of the expression of *LEF-1* and *TCF-1* transcription factors, stem cell markers *LGR5* and *NANOG* (E), and *CCND1, MMP7* and *C-MYC* (F). **(G)** Protein expression levels of E-cadherin (differentiation marker), LGR5, Hakai and NANOG (stem cell markers) upon treatment with Hakin-1 (50 µM) were analysed by Western blot, using the indicated antibodies. Vinculin was used as loading control. Protein bands were quantified using ImageJ and normalized to loading control. Results are expressed as mean ± SEM. T-test was performed for statistical analysis (*p < 0.05; **p < 0.01; ***p < 0.001).

Given the observed effects on stemness, we next analysed the potential impact of Hakin-1 on the subcellular localization of LGR5 and MUC2 in tumourspheres. Moreover, we examined the effect of Hakai inhibition on E-cadherin and LRP4 localization in tumourspheres using immunofluorescence, given E-cadherin’s role in EMT and LRP4 as a novel Hakai-interacting protein linked to Wnt/β-catenin signalling. Hakin-1 treatment modestly increased E-cadherin levels at cell-cell contacts (¡Error! No se encuentra el origen de la referencia.**A**). In contrast, Hakin-1 treatment resulted in a reduction in LGR5 levels (¡Error! No se encuentra el origen de la referencia.**B**), suggesting a potential decrease in the stemness of cancer cells. On the other hand, Hakin-1 treatment increased the expression of the differentiation marker MUC2 (¡Error! No se encuentra el origen de la referencia.**D**). As expected, LRP4, as a target for Hakai, showed increased expression levels upon Hakin-1 treatment (¡Error! No se encuentra el origen de la referencia.**C**). Taken together our results suggest that Hakin-1 impairs tumoursphere formation by reducing stemness-associated features, disrupting 3D structural integrity, and promoting cellular differentiation.

In conclusion, our findings identify Hakai as a critical regulator of Wnt/β-catenin signalling in colorectal cancer as illustrated by the model in **Fig. 7**. Hakai ubiquitinates LRP4, targeting it for proteasomal degradation. The degradation of membrane-bound LRP4 may relieve its inhibitory effect on Wnt signalling, thereby facilitating Hakai-mediated cooperation with LRP6 to potentiate pathway activation. Consequently, promoting the expression of Wnt/β-catenin target genes that induce stem-like properties in colon cancer. Pharmacological inhibition of Hakai may reduce stemness-associated features and promote cellular differentiation in colorectal cancer cells.

## Discussion

In this study, we provide evidence that the E3 ubiquitin-ligase Hakai plays a pivotal role in regulating CSC properties in colorectal cancer. Using colon cancer tumoursphere as a model of CSC, we show that Hakai modulates self-renewal, differentiation and Wnt/β-catenin signalling. Silencing Hakai significantly impairs self-renewal capacity, as shown by a marked reduction in tumoursphere number and size (**Fig. 1B–D**), and leads to downregulation of key CSC markers, including the transcriptional Wnt target gene *LGR5* (**Fig. 1F**) [7,36]. These results are consistent with our previous findings showing that tumourspheres derived from CRC cells exhibit increased *CBLL1* (Hakai gene) and *LGR5* mRNA levels compared to monolayer cultures [29]. Our mechanistic studies suggest that Hakai could promote the acquisition of CSC properties via the hyperactivation of the Wnt/β-catenin signalling pathway in colon cancer. We further identified the negative regulator of Wnt/β-catenin signalling, LRP4, as a novel substrate of Hakai-mediated ubiquitination. Given our biochemical studies and the important role of Wnt signalling in the development of CSCs, we hypothesize that Hakai’s effect on stemness is at least partly through the newly identified substrate LRP4. Notably, this is the first study to implicate LRP4 in Wnt signalling within colorectal CSCs, expanding its role beyond its previously recognized function as a negative regulator during developmental processes. Hakai overexpression decreased LRP4 protein levels and its surface localization (**Fig. 3A-B**), while Hakai depletion led to its accumulation. Co-immunoprecipitation assays confirmed the interaction between Hakai and LRP4, and ubiquitination studies demonstrated that Hakai targets LRP4 for proteasomal degradation (**Fig. 3C-E**). These findings identify LRP4 as a novel substrate of Hakai and suggest that Hakai contributes to Wnt/β-catenin pathway hyperactivation in CRC by removing LRP4-mediated inhibition of LRP6. According to this, a recent study demonstrates that although WNT-induced FZD-LRP6 interaction occurs, this is not sufficient to initiate downstream Wnt/β-catenin signalling, indicating that additional regulatory mechanisms are required for full pathway activation [37]. Moreover, the co-expression of *CBLL1* and *LGR5* was proposed for the identification of a subpopulation of CMS2 patients with more aggressive biological characteristics [29]. Notably, this may include cooperation with LRP6 to overcome LRP4-mediated inhibition, a mechanism consistent with previous reports of LRP4 antagonizing LRP6 signalling, especially in developmental contexts [9,10]. We further evaluated the impact of pharmacological inhibition of Hakai using Hakin-1, a compound targeting the HYB domain responsible for its ubiquitin-ligase activity [22]. Treatment with Hakin-1 significantly reduced tumoursphere formation and stem cell marker expression (**Fig. 5C-G**), while increasing differentiation markers such as MUC2, which is indicative of goblet cell lineage commitment [38]. Importantly, Hakin-1 also exerted effects on pre-established tumourspheres, demonstrating its potential in both preventive and therapeutic contexts (**Fig. 5, Fig. S1-S3**). These results were further corroborated by increased E-cadherin and LRP4 expression (**Fig. 6**), suggesting that Hakin-1 inhibits EMT and the hyperactivation of Wnt/β-catenin signalling.

**Fig 6.**
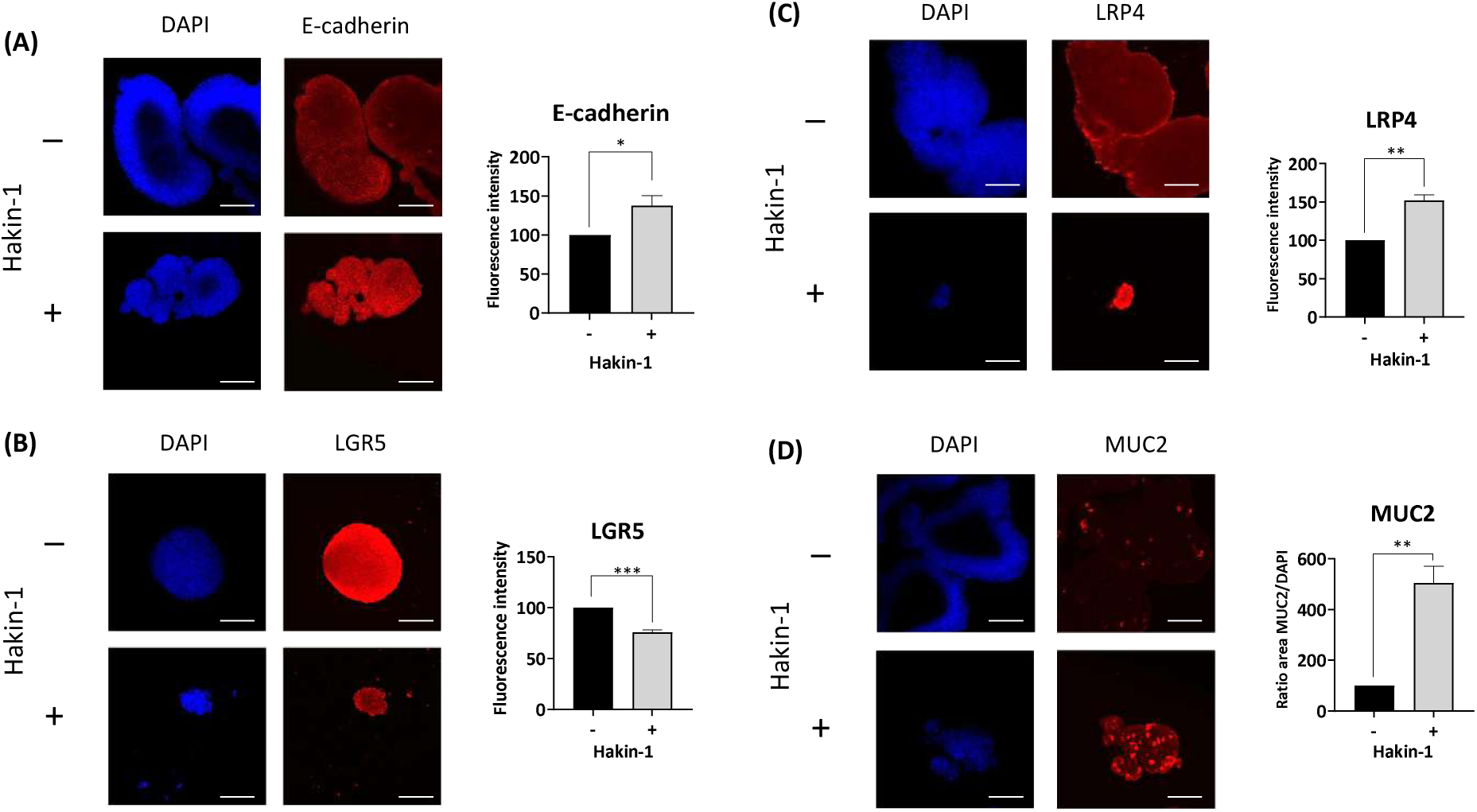
Localization of stem cell and differentiation markers in HT29 tumourspheres upon Hakin-1 treatment. Immunofluorescence analysis of HT29 tumourspheres treated with DMSO (vehicle control) or Hakin-1 (50 µM). **(A–D)** Left panels: representative immunofluorescence images of E-cadherin (A), LGR5 (B), LRP4 (C), and MUC2 (D). Images were captured using a 20x objective (scale bar = 50 µm); magnified images (circle images) were generated by digital zoom. Right panels: quantitative analysis of marker expression. For E-cadherin, LRP4, and LGR5, fluorescence intensity was normalized to area and quantified across at least 5 tumourspheres per condition, shown as scatter plots. For MUC2, the number of positive cells was quantified by measuring the stained area and normalizing to DAPI-stained nuclear area. Data are presented as mean ± SEM from three independent experiments. Statistical significance was calculated using unpaired *t*-tests (*p < 0.05, **p < 0.01, ***p < 0.001) in GraphPad Prism.

**Fig 7.**
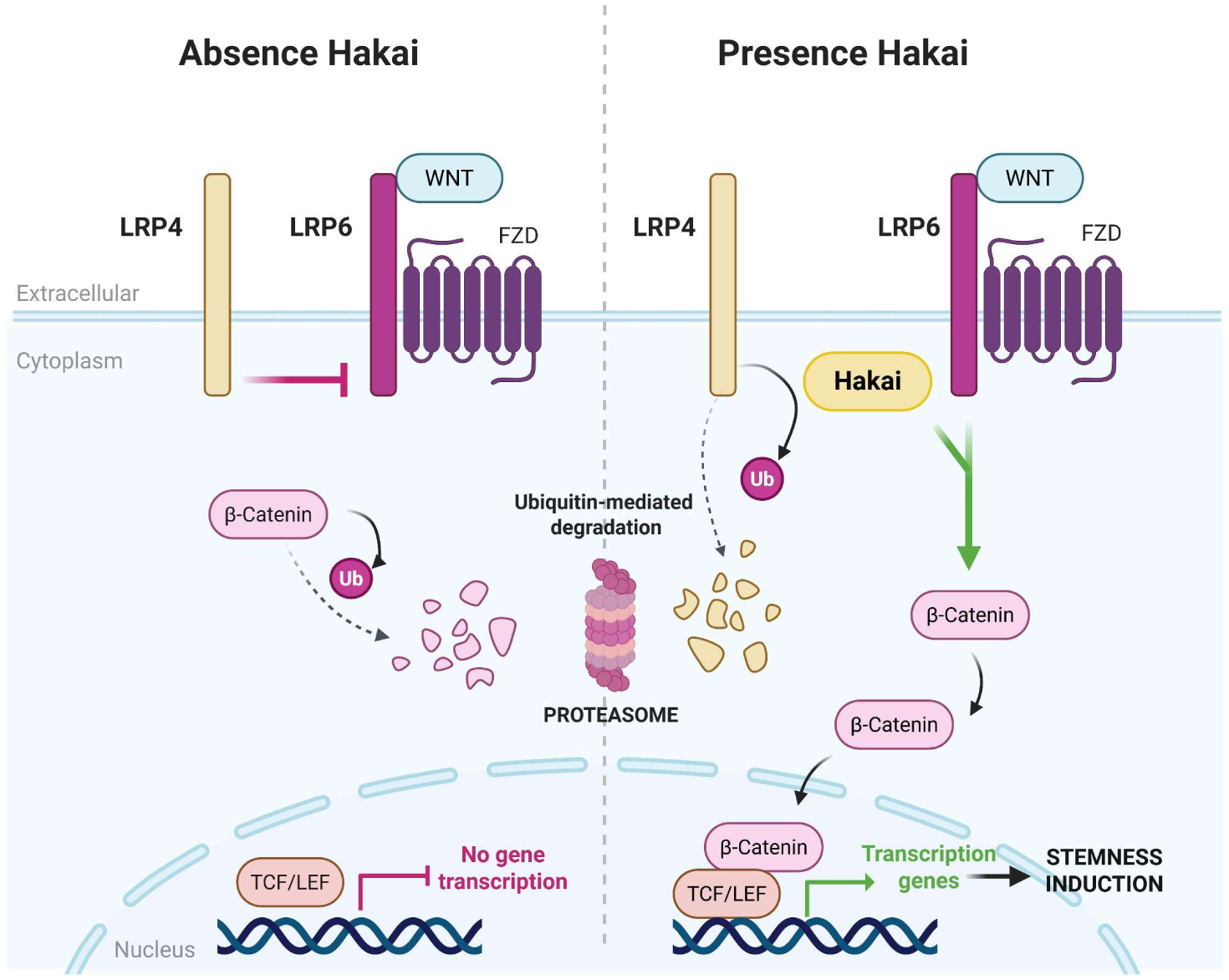
Model proposed for Hakai action in Wnt/β-Catenin signalling pathway. Hakai absence: LRP4 acts as an antagonist at the cell membrane, inhibiting Wnt/β-catenin signalling by interfering with LRP6 activity. As a result, cytoplasmic β-catenin is targeted for proteasomal degradation, and Wnt target genes remain transcriptionally repressed. **In the presence of Hakai**, ubiquitination of LRP4 is induced, thereby promoting its degradation. This action of Hakai can overcome the inhibitory effect of LRP4, resulting in increased LRP6-mediated Wnt signalling. The increased Wnt/β-catenin signalling may promote stem cell properties in tumourspheres. Image created with BioRender.com.

The interplay between Hakai and Wnt/β-catenin signalling refines our understanding of CSC regulation. While mutations in *APC*, *CTNNB1*, or *AXIN* frequently lead to Wnt/β-catenin pathway activation in CRC [3,39] our findings suggest that such genetic alterations are not sufficient to induce a full CSC phenotype. Instead, CSCs may require additional signals or co-factors to enhance Wnt/β-catenin signalling activity. Only cells with the highest nuclear β-catenin activity exhibit CSC traits [5], highlighting the importance of post-translational regulators like Hakai in enhancing Wnt/β-catenin pathway. Indeed, multiple E3 ubiquitin-ligases modulate Wnt signalling through degradation of pathway components. For example, Mindbomb 1 enhances Wnt/β-catenin activity by targeting the non-canonical receptor RYK [40], while Itch ubiquitinates LRP6 to promote endocytosis and downstream signalling [41]. Conversely, RNF43 and ZNRF3 negatively regulate Wnt/β-catenin signalling by promoting the ubiquitination and internalization of Frizzled receptors, thus attenuating pathway activation [42,43]. Compared to these ligases, Hakai appears to occupy a unique niche by simultaneously targeting both E-cadherin and LRP4, thereby integrating EMT and Wnt/β-catenin signalling into a common regulatory axis. In addition to modulating both signalling pathways, Hakai may also influence metabolic adaptations essential for CSC maintenance. Proteomic profiling revealed that Hakai silencing leads to a marked reduction in PRPS2 (**Fig. S4**), a purine biosynthesis enzyme regulated by MYC signalling and previously implicated in supporting CSC survival and proliferation [44,45]. Given that *C-MYC* was the first gene recognized as a target of the Wnt/β-catenin pathway in CRC [46], this finding suggests that Hakai might promote metabolic activity indirectly through its impact on Wnt-driven *C-MYC* expression. These results are consistent with our previous results, which demonstrated an association between *CBLL1* expression and the CMS2 subtype in CRC patients, also characterized by Myc pathway activation [29]. This opens the possibility that Hakai may exert a metabolic influence that complements its known effects on signalling and differentiation, suggesting a new direction for future investigation.

Mechanistically, our findings suggest that Hakai not only facilitates Wnt/β-catenin signalling by promoting the degradation of LRP4, but may also contribute to LRP6-mediated pathway activation. Reporter assays under WNT3A stimulation showed that co-expression of Hakai and LRP6 restored Wnt/β-catenin activity suppressed by LRP4 more effectively than either factor alone (**Fig. 4B**). These observations imply that Hakai helps counteract LRP4’s inhibitory role, possibly by promoting the clearance of suppressive inputs and thereby facilitating the assembly or function of activating complexes. This interpretation is consistent with previous reports describing the antagonistic role of LRP4 in modulating LRP6-mediated Wnt/β-catenin activation, particularly in developmental systems such as limb formation, bone development, and the neuromuscular junction [9,10,13,47,48], and aligns with models of Wnt/β-catenin signalling that emphasize the importance of the changing interactions and arrangements of receptors on the cell surface [49].

Hakin-1’s therapeutic profile compares favourably with other agents targeting the Wnt/β-catenin pathway. For instance, anti-LRP6 nanobodies and R-spondin 3 (RSPO3) inhibitors reduce tumour growth and induce differentiation by exhausting the CSC pool [50,51] Similarly, APC restoration in inducible mouse models similarly drives differentiation and suppresses tumour relapse [52] Like these agents, Hakin-1 promotes MUC2 expression and downregulates LGR5, suggesting a robust differentiation response. Notably, unlike broad Wnt inhibitors, Hakin-1 has not been associated with systemic toxicity, based on previous histopathological analyses in treated animals [22] However, the *in vivo* impact of Hakai’s pharmacological inhibition on the Wnt/β-catenin signalling cascade remains to be elucidated. Targeting E3 ligases for cancer therapy is gaining traction due to their substrate specificity and regulatory capacity. While bortezomib and carfilzomib target the proteasome broadly, leading to side effects such as neuropathy and myelosuppression [53,54], E3 ligase inhibitors like MLN4924, which targets the NEDD8-activating enzyme to inhibit Cullin-RING ligases, and Hakin-1, demonstrate greater substrate specificity and improved lower toxicity[55,56]. Our data show that Hakin-1 does not impair apoptosis but instead shifts tumour cells toward differentiation, a therapeutic endpoint now increasingly recognized as essential for long-term tumour control. Notably, Hakai-silencing shows a more pronounced effect on cell proliferation compared to the more limited impact of Hakin-1 targeting Hakai’s E3 ligase activity. This is consistent with previous reports showing that full knockdown of Hakai markedly inhibits proliferation and decreases Cyclin D1 protein levels [22,34]. In contrast, Hakin-1 treatment shows only a modest effect in proliferation *in vitro* and *in vivo* [22] and does not significantly alter proliferation-associated targets such as *C-MYC* or *CCND1* (**Fig. 5F**). These findings support the hypothesis that other functional domains of Hakai may be responsible for its effects on proliferation, as the inhibition of the HYB domain, responsible for its E3 ubiquitin-ligase activity, shows only a limited impact when targeted alone. From a translational perspective, the ability of Hakin-1 to reduce stemness, enhance differentiation, and interfere with both Wnt/β-catenin signalling and EMT underscores its value as a therapeutic agent capable of targeting key mechanisms underlying cancer resistance and metastasis. Its impact on both the initiation and maintenance of tumourspheres, together with its favourable safety profile, positions it as a strong candidate for combinatorial regimens targeting CSCs alongside standard chemotherapy.

## Conclusions

In conclusion, our findings define a new functional role for Hakai in colorectal cancer, highlighting its involvement in CSC regulation and Wnt/β-catenin signalling through LRP4-mediated modulation. The development of Hakin-1 provides a promising therapeutic strategy to target CSCs by inhibiting Hakai-mediated ubiquitination, enhancing differentiation, and attenuating Wnt/β-catenin pathway activity. These insights support further investigation of Hakai inhibitors as components of CSC-directed therapies in colorectal cancer and offer a rational approach for improving long-term treatment outcomes.

## Glossary

3-MA: 3-Methyladenine
AKT: Protein kinase B
APC: Adenomatous polyposis coli
APS: Ammonium persulfate
ATCC: American Type Culture Collection
b-hFGF: Basic human fibroblast growth factor
BCA: Bicinchoninic acid assay
BMP: Bone morphogenetic protein
BSA: Bovine serum albumin
C19ORF43: Telomerase RNA component interacting RNase
CBLL1: Casitas B-lineage lymphoma like-1, Hakai gene
CCND1: Cyclin D1 gene
CMS: Consensus Molecular Subtype
CRC: Colorectal cancer
CSCs: Cancer Stem Cells
CT: Threshold cycle
DAPI: 4’,6-diamidino-2-phenylindole
DDA: Data-dependent acquisition
DKK: Dickkopf
DMEM: Dulbecco’s Modified Eagle’s Medium
DMSO: Dimethyl sulfoxide
Dox: Doxycycline
DVL: Dishevelled
EGF: Epidermal growth factor
EMT: Epithelial-mesenchymal transition
ESI: Electrospray ionization
FA: Formic acid
FBS: Fetal bovine serum
FZD: Frizzled transmembrane receptors
GAPDH: Glyceraldehyde 3-phosphate dehydrogenase
GSK3β: Glycogen synthase kinase-3 β
Hakin-1: Hakai inhibitor 1
HRP: Horseradish peroxidase
HYB: Hakai-pY-binding
IAA: Iodoacetamide
LC-MS/MS: Liquid chromatography with tandem mass spectrometry
LFQ: Label-free quantification
LGR5: Leucine-rich repeat-containing G-protein coupled Receptor 5
LRP4: Low density lipoprotein receptor-related protein 4
LRP5: Low density lipoprotein receptor-related protein 5
LRP6: Low density lipoprotein receptor-related protein 6
MESD: Mesoderm development LRP chaperone
MNAR: Missing not at random
MUC2: Mucin2
MYC: MYC proto-oncogene
NIT1: Nitrilase homolog 1
PASEF: Parallel accumulation–serial fragmentation
PBL: Passive lysis buffer
PBS: Phosphate-buffered saline
PBST: Phosphate-buffered saline, 0.1% Triton X-100
PCA: Principal component analysis
PCs: Principal components
PCR: Polymerase chain reaction
PFA: Paraformaldehyde
PI3K: Phosphatidylinositol 3-kinase
PMSF: Phenylmethylsulfonyl fluoride
PRPS2: Phosphoribosyl pyrophosphate synthetase 2
pTyr: Tyrosine residue phosphorylation, or phosphotyrosine
PVDF: Polyvinylidene difluoride
RABGGTA: Rab geranylgeranyltransferase alpha subunit
RNF43: Ring finger protein 43
RSPO: R-spondin
RT: Room temperature
SLC35F2: Solute carrier family 35 member F2 SOST Sclerostin
TBS-T: Tris buffered saline-Tween 20
TCF/LEF: T-cell factor/lymphoid enhancer-binding factor
TimsTOF: Trapped ion mobility spectrometry quadrupole time-of-flight
UBXN7: UBX domain-containing protein 7
UFSP2: Ufm1-specific protease 2
WDR26: WD repeat-containing protein 26
WISE: Wnt inhibitory secreted protein
ZNRF3: Zinc and ring finger protein 3

## Declarations

### Availability of data and materials

All data generated or analysed during this study are included in this published article and its supplementary information files. The datasets used and/or analysed during the current study are available from the corresponding author on reasonable request.

### Competing interests

A.F, A.R.A and L.J. are inventors on patents related to inhibitors of the E3 ubiquitin-ligase Hakai.

### Funding sources

This study has been funded by the Instituto de Salud Carlos III (ISCIII) through the project numbers PI21/00238 and FORT23/00010, and co-funded by the European Union. The project that gave rise to these results has received funding from “la Caixa” Foundation and the European Institute of Innovation and Technology, EIT (body of the European Union that receives support from the European Union’s Horizon 2020 research and innovation program), under the grant agreement LCF/TR/CC21/52490003 and is also supported by Consolidation of Competitive Research (IN607B 2023/12) from Agencia Gallega de Innovación (GAIN) from Xunta de Galicia. This work was supported by a grant from the Scientific Foundation of the Spanish Association against Cancer (INNOV235141FIGU). A.R.A. and G.A. are supported by a predoctoral contract (PRDLC21591RODR and PRDLC234251ALFO, respectively) from Fundación Científica Asociación Española Contra el Cáncer (AECC/AECC) and L.J. is supported by FPU contract (FPU021/05350) from Ministerio de Universidades (Spain).

### Authors’ contributions

Conceptualization: A.F.; investigation and methodology: A.R.-A., L.J., I.J.; data curation and formal analysis: A.R.-A.; bioinformatic analysis: G.A.; writing—original draft: A.F. and A.R.-A; writing—review & editing: A.F., A.R.-A., I.J., M.M.; with contributions from all authors; supervision, project administration, and funding acquisition: A.F. All authors have read and agreed to the published version of the manuscript

## Acknowledgements

We thank Dr. Yasuyuki Fujita (Hokkaido University, Japan) for generously providing the pcDNA-FLAG-Hakai and pBSSR-HA-Ubiquitin constructs, and Dr. Tatsuo Suzuki (Shinshu University School of Medicine, Japan) for kindly providing the pcDNA-LRP4 plasmid. We would like to thank the Proteomics Unit 30 of NANBIOSIS (CIBER-BBN) at the INIBIC for the support with the sample preparation and the Proteomics laboratory at the Interdisciplinary Center for Chemistry and Biology (CICA), UDC for their expertise with the LC-MS/MS and data analysis

## APPENDIX A. Supplemental information

**Table S1.**
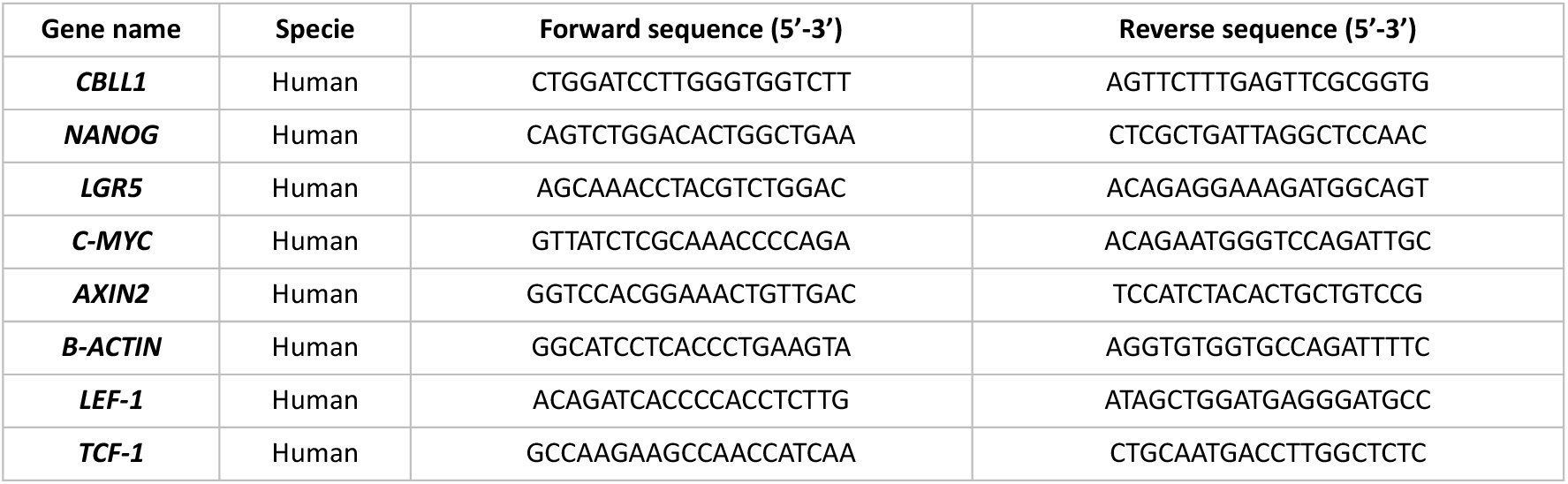
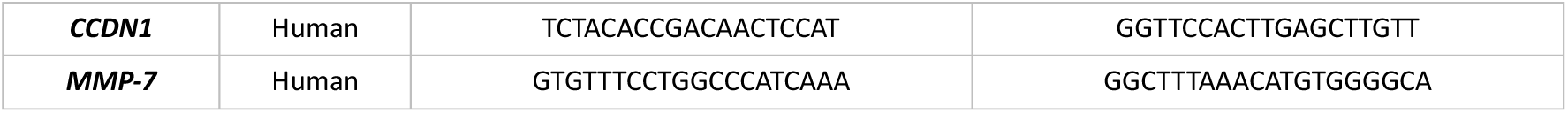
Primers sequences used for RT-qPCR. Sequences of the forward and reverse primers used for quantitative reverse transcription PCR (RT-qPCR) to amplify target genes. Primer sequences were designed using BLAST and Primer3. Gene names correspond to those used in the main text.

**Table S2.** Predicted Hakai substrate interactions by bioinformatics. UbiBrowser platform was used to determine the predicted Hakai-substrate interaction network. A total of 116 proteins were found as potential predicted substrates for Hakai-mediated ubiquitination [32].

|  | SwissProt ID (E3) | Gene Symbol (E3) | SwissProt ID (Substrate) | Gene Symbol (Substrate) | Confidence Score | Species |
| --- | --- | --- | --- | --- | --- | --- |
| 1 | Q75N03 | CBLL1 | O00499 | BIN1 | 0.851 | H.sapiens |
| 2 | Q75N03 | CBLL1 | O15230 | LAMA5 | 0.851 | H.sapiens |
| 3 | Q75N03 | CBLL1 | O15353 | FOXN1 | 0.851 | H.sapiens |
| 4 | Q75N03 | CBLL1 | O75096 | LRP4 | 0.851 | H.sapiens |
| 5 | Q75N03 | CBLL1 | O75147 | OBSL1 | 0.851 | H.sapiens |
| 6 | Q75N03 | CBLL1 | O75473 | LGR5 | 0.851 | H.sapiens |
| 7 | Q75N03 | CBLL1 | P01133 | EGF | 0.851 | H.sapiens |
| 8 | Q75N03 | CBLL1 | P02751 | FN1 | 0.851 | H.sapiens |
| 9 | Q75N03 | CBLL1 | P07996 | THBS1 | 0.851 | H.sapiens |
| 10 | Q75N03 | CBLL1 | P08631 | HCK | 0.851 | H.sapiens |
| 11 | Q75N03 | CBLL1 | P08922 | ROS1 | 0.851 | H.sapiens |
| 12 | Q75N03 | CBLL1 | P08F94 | PKHD1 | 0.851 | H.sapiens |
| 13 | Q75N03 | CBLL1 | P11362 | FGFR1 | 0.851 | H.sapiens |
| 14 | Q75N03 | CBLL1 | P13688 | CEACAM1 | 0.851 | H.sapiens |
| 15 | Q75N03 | CBLL1 | P15391 | CD19 | 0.851 | H.sapiens |
| 16 | Q75N03 | CBLL1 | P17948 | FLT1 | 0.851 | H.sapiens |
| 17 | Q75N03 | CBLL1 | P23458 | JAK1 | 0.851 | H.sapiens |
| 18 | Q75N03 | CBLL1 | P23634 | ATP2B4 | 0.851 | H.sapiens |
| 19 | Q75N03 | CBLL1 | P25391 | LAMA1 | 0.851 | H.sapiens |
| 20 | Q75N03 | CBLL1 | P28827 | PTPRM | 0.851 | H.sapiens |
| 21 | Q75N03 | CBLL1 | P31276 | HOXC13 | 0.851 | H.sapiens |
| 22 | Q75N03 | CBLL1 | P36888 | FLT3 | 0.851 | H.sapiens |
| 23 | Q75N03 | CBLL1 | P42229 | STAT5A | 0.851 | H.sapiens |
| 24 | Q75N03 | CBLL1 | P43405 | SYK | 0.851 | H.sapiens |
| 25 | Q75N03 | CBLL1 | P49286 | MTNR1B | 0.851 | H.sapiens |
| 26 | Q75N03 | CBLL1 | P51608 | MECP2 | 0.851 | H.sapiens |
| 27 | Q75N03 | CBLL1 | P51692 | STAT5B | 0.851 | H.sapiens |
| 28 | Q75N03 | CBLL1 | P52179 | MYOM1 | 0.851 | H.sapiens |
| 29 | Q75N03 | CBLL1 | P52333 | JAK3 | 0.851 | H.sapiens |
| 30 | Q75N03 | CBLL1 | P52735 | VAV2 | 0.851 | H.sapiens |
| 31 | Q75N03 | CBLL1 | P54296 | MYOM2 | 0.851 | H.sapiens |
| 32 | Q75N03 | CBLL1 | P54762 | EPHB1 | 0.851 | H.sapiens |
| 33 | Q75N03 | CBLL1 | P56539 | CAV3 | 0.851 | H.sapiens |
| 34 | Q75N03 | CBLL1 | Q07889 | SOS1 | 0.851 | H.sapiens |
| 35 | Q75N03 | CBLL1 | Q13023 | AKAP6 | 0.851 | H.sapiens |
| 36 | Q75N03 | CBLL1 | Q16288 | NTRK3 | 0.851 | H.sapiens |
| 37 | Q75N03 | CBLL1 | Q16620 | NTRK2 | 0.851 | H.sapiens |
| 38 | Q75N03 | CBLL1 | Q5VTT5 | MYOM3 | 0.851 | H.sapiens |
| 39 | Q75N03 | CBLL1 | Q6Q0C0 | TRAF7 | 0.851 | H.sapiens |
| 40 | Q75N03 | CBLL1 | Q8WU20 | FRS2 | 0.851 | H.sapiens |
| 41 | Q75N03 | CBLL1 | Q8WZ42 | TTN | 0.851 | H.sapiens |
| 42 | Q75N03 | CBLL1 | Q96BA8 | CREB3L1 | 0.851 | H.sapiens |
| 43 | Q75N03 | CBLL1 | Q99996 | AKAP9 | 0.851 | H.sapiens |
| 44 | Q75N03 | CBLL1 | Q9BXB1 | LGR4 | 0.851 | H.sapiens |
| 45 | Q75N03 | CBLL1 | Q9HAZ2 | PRDM16 | 0.851 | H.sapiens |
| 46 | Q75N03 | CBLL1 | Q9HC29 | NOD2 | 0.851 | H.sapiens |
| 47 | Q75N03 | CBLL1 | Q9NRA0 | SPHK2 | 0.851 | H.sapiens |
| 48 | Q75N03 | CBLL1 | Q9NYQ6 | CELSR1 | 0.851 | H.sapiens |
| 49 | Q75N03 | CBLL1 | Q9UIU6 | SIX4 | 0.851 | H.sapiens |
| 50 | Q75N03 | CBLL1 | Q9UKW4 | VAV3 | 0.851 | H.sapiens |
| 51 | Q75N03 | CBLL1 | P00533 | EGFR | 0.850 | H.sapiens |
| 52 | Q75N03 | CBLL1 | P01127 | PDGFB | 0.850 | H.sapiens |
| 53 | Q75N03 | CBLL1 | P02671 | FGA | 0.850 | H.sapiens |
| 54 | Q75N03 | CBLL1 | P02675 | FGB | 0.850 | H.sapiens |
| 55 | Q75N03 | CBLL1 | P05019 | IGF1 | 0.850 | H.sapiens |
| 56 | Q75N03 | CBLL1 | P06241 | FYN | 0.850 | H.sapiens |
| 57 | Q75N03 | CBLL1 | P07333 | CSF1R | 0.850 | H.sapiens |
| 58 | Q75N03 | CBLL1 | P07359 | GP1BA | 0.850 | H.sapiens |
| 59 | Q75N03 | CBLL1 | P07947 | YES1 | 0.850 | H.sapiens |
| 60 | Q75N03 | CBLL1 | P08514 | ITGA2B | 0.850 | H.sapiens |
| 61 | Q75N03 | CBLL1 | P09327 | VIL1 | 0.850 | H.sapiens |
| 62 | Q75N03 | CBLL1 | P10721 | KIT | 0.850 | H.sapiens |
| 63 | Q75N03 | CBLL1 | P16066 | NPR1 | 0.850 | H.sapiens |
| 64 | Q75N03 | CBLL1 | P18206 | VCL | 0.850 | H.sapiens |
| 65 | Q75N03 | CBLL1 | P21333 | FLNA | 0.850 | H.sapiens |
| 66 | Q75N03 | CBLL1 | P21802 | FGFR2 | 0.850 | H.sapiens |
| 67 | Q75N03 | CBLL1 | P22607 | FGFR3 | 0.850 | H.sapiens |
| 68 | Q75N03 | CBLL1 | P29322 | EPHA8 | 0.850 | H.sapiens |
| 69 | Q75N03 | CBLL1 | P29350 | PTPN6 | 0.850 | H.sapiens |
| 70 | Q75N03 | CBLL1 | P33151 | CDH5 | 0.850 | H.sapiens |
| 71 | Q75N03 | CBLL1 | P35367 | HRH1 | 0.850 | H.sapiens |
| 72 | Q75N03 | CBLL1 | P40189 | IL6ST | 0.850 | H.sapiens |
| 73 | Q75N03 | CBLL1 | P42338 | PIK3CB | 0.850 | H.sapiens |
| 74 | Q75N03 | CBLL1 | P49747 | COMP | 0.850 | H.sapiens |
| 75 | Q75N03 | CBLL1 | P49767 | VEGFC | 0.850 | H.sapiens |
| 76 | Q75N03 | CBLL1 | Q02763 | TEK | 0.850 | H.sapiens |
| 77 | Q75N03 | CBLL1 | Q06418 | TYRO3 | 0.850 | H.sapiens |
| 78 | Q75N03 | CBLL1 | Q12979 | ABR | 0.850 | H.sapiens |
| 79 | Q75N03 | CBLL1 | Q5VY43 | PEAR1 | 0.850 | H.sapiens |
| 80 | Q75N03 | CBLL1 | Q68CJ9 | CREB3L3 | 0.850 | H.sapiens |
| 81 | Q75N03 | CBLL1 | Q8N9M5 | TMEM102 | 0.850 | H.sapiens |
| 82 | Q75N03 | CBLL1 | Q96QS1 | TSPAN32 | 0.850 | H.sapiens |
| 83 | Q75N03 | CBLL1 | Q99650 | OSMR | 0.850 | H.sapiens |
| 84 | Q75N03 | CBLL1 | Q9BRC7 | PLCD4 | 0.850 | H.sapiens |
| 85 | Q75N03 | CBLL1 | Q9UL54 | TAOK2 | 0.850 | H.sapiens |
| 86 | Q75N03 | CBLL1 | P17813 | ENG | 0.822 | H.sapiens |
| 87 | Q75N03 | CBLL1 | P35580 | MYH10 | 0.822 | H.sapiens |
| 88 | Q75N03 | CBLL1 | P36402 | TCF7 | 0.822 | H.sapiens |
| 89 | Q75N03 | CBLL1 | Q03395 | ROM1 | 0.822 | H.sapiens |
| 90 | Q75N03 | CBLL1 | Q12809 | KCNH2 | 0.822 | H.sapiens |
| 91 | Q75N03 | CBLL1 | Q13009 | TIAM1 | 0.822 | H.sapiens |
| 92 | Q75N03 | CBLL1 | Q6KC79 | NIPBL | 0.822 | H.sapiens |
| 93 | Q75N03 | CBLL1 | Q6UWI4 | SHISA2 | 0.822 | H.sapiens |
| 94 | Q75N03 | CBLL1 | Q9UJU2 | LEF1 | 0.822 | H.sapiens |
| 95 | Q75N03 | CBLL1 | Q9UNE0 | EDAR | 0.822 | H.sapiens |
| 96 | Q75N03 | CBLL1 | Q9Y264 | ANGPT4 | 0.822 | H.sapiens |
| 97 | Q75N03 | CBLL1 | O75038 | PLCH2 | 0.821 | H.sapiens |
| 98 | Q75N03 | CBLL1 | P20827 | EFNA1 | 0.821 | H.sapiens |
| 99 | Q75N03 | CBLL1 | P35579 | MYH9 | 0.821 | H.sapiens |
| 100 | Q75N03 | CBLL1 | Q01826 | SATB1 | 0.821 | H.sapiens |
| 101 | Q75N03 | CBLL1 | Q15173 | PPP2R5B | 0.821 | H.sapiens |
| 102 | Q75N03 | CBLL1 | Q6ZWH5 | NEK10 | 0.821 | H.sapiens |
| 103 | Q75N03 | CBLL1 | P07948 | LYN | 0.810 | H.sapiens |
| 104 | Q75N03 | CBLL1 | P09619 | PDGFRB | 0.810 | H.sapiens |
| 105 | Q75N03 | CBLL1 | P16234 | PDGFRA | 0.810 | H.sapiens |
| 106 | Q75N03 | CBLL1 | Q7Z6A9 | BTLA | 0.810 | H.sapiens |
| 107 | Q75N03 | CBLL1 | O75593 | FOXH1 | 0.809 | H.sapiens |
| 108 | Q75N03 | CBLL1 | O96020 | CCNE2 | 0.809 | H.sapiens |
| 109 | Q75N03 | CBLL1 | P24864 | CCNE1 | 0.809 | H.sapiens |
| 110 | Q75N03 | CBLL1 | P25445 | FAS | 0.809 | H.sapiens |
| 111 | Q75N03 | CBLL1 | Q13485 | SMAD4 | 0.809 | H.sapiens |
| 112 | Q75N03 | CBLL1 | Q15431 | SYCP1 | 0.809 | H.sapiens |
| 113 | Q75N03 | CBLL1 | Q16181 | SEPTIN7 | 0.809 | H.sapiens |
| 114 | Q75N03 | CBLL1 | Q86VD1 | MORC1 | 0.809 | H.sapiens |
| 115 | Q75N03 | CBLL1 | Q9UH03 | SEPTIN3 | 0.809 | H.sapiens |
| 116 | Q75N03 | CBLL1 | Q9Y615 | ACTL7A | 0.809 | H.sapiens |

## Supplementary figures

**Fig. S1.**
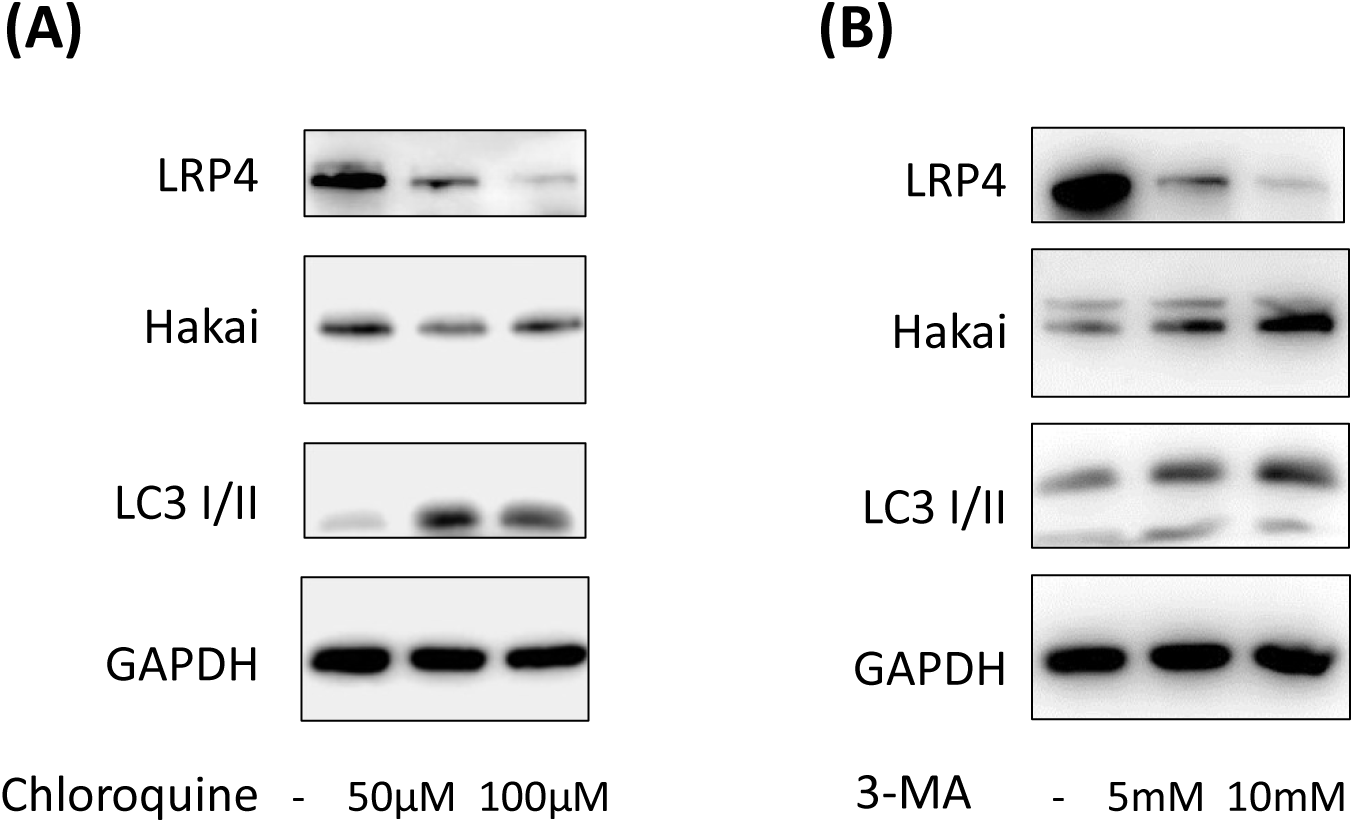
LRP4 protein level is not recovered in presence of lysosome inhibitor Chloroquine nor in presence of autophagy inhibitor 3-MA. Endogenous LRP4 and Hakai levels in HCT116 cells treated with (A) lysosome inhibitor Chloroquine and (B) autophagy inhibitor 3-MA analyzed by Western blot. LC3 I/II was used as a positive control for chloroquine and 3-MA treatment and GAPDH as a loading control.

**Fig. S2.**
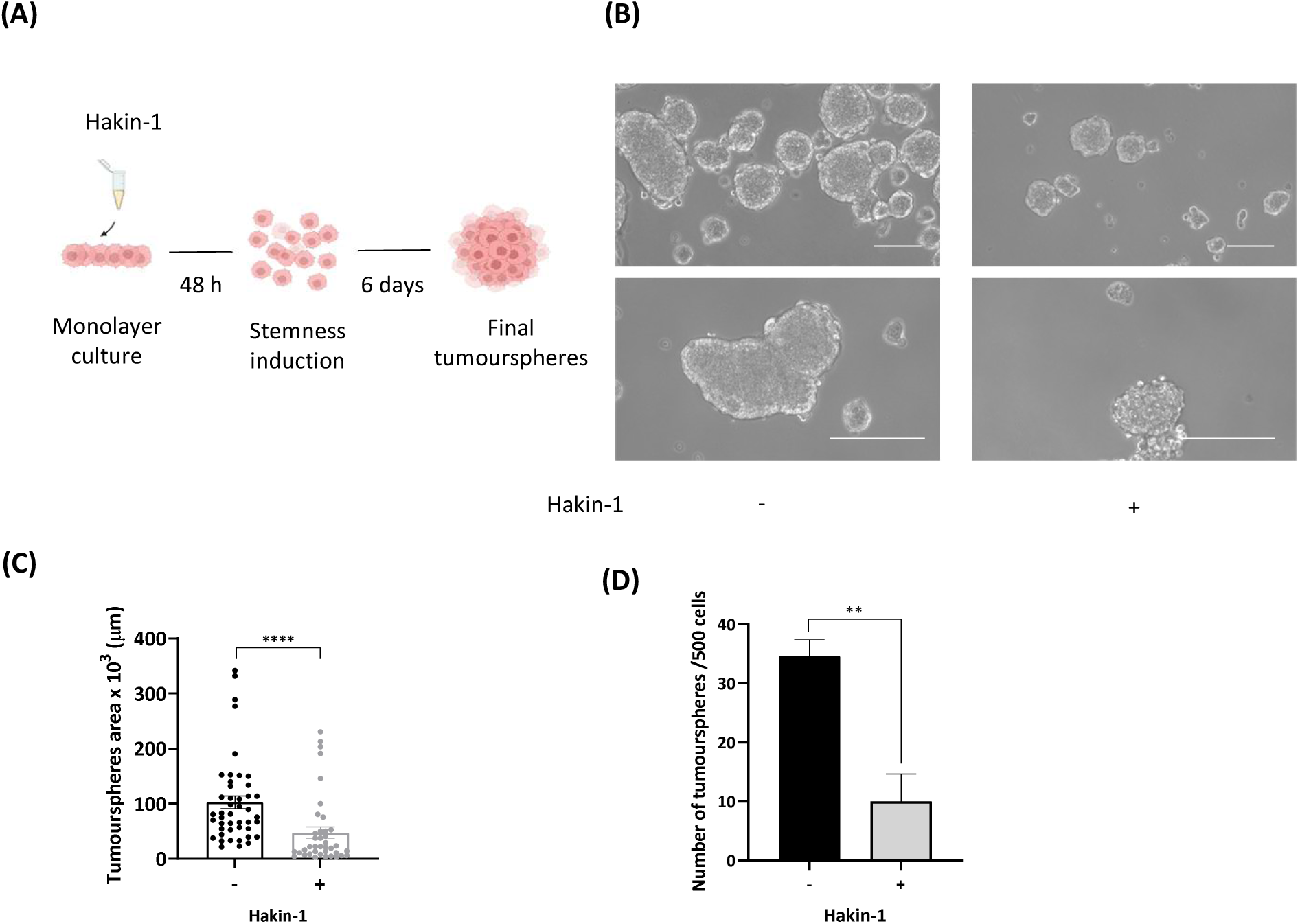
Impact of the effect of Hakin-1 prior to stemness induction. **(A)** Schematic workflow. Hakin-1 treatment was administered 48 hours prior to the induction of tumoursphere formation. **(B)** Representative images of tumourspheres in HT29 treated with DMSO (control) or 50 μM Hakin-1 treatment on day 6 after stemness condition. Images were captured using a 4x (upper images, scale bar: 250 μm) or 10x (lower images, scale bar: 200 μm) objective. **(C)** Tumoursphere size measurement. Images of tumourspheres derived from HT29 cells treated with DMSO (control) or Hakin-1 were captured, and surface area quantification was performed using ImageJ software. Tumoursphere area of at least 15 tumourspheres per experiment were measured. **(D)** Tumourspheres formation quantification. The quantification of tumourspheres formed by HT29 cells treated with DMSO (control) or Hakin-1 was performed by self-renewal assay. Statistical analysis was performed using a t-test, with significance levels as *p < 0.05 and ****p < 0.0001.

**Fig. S3.**
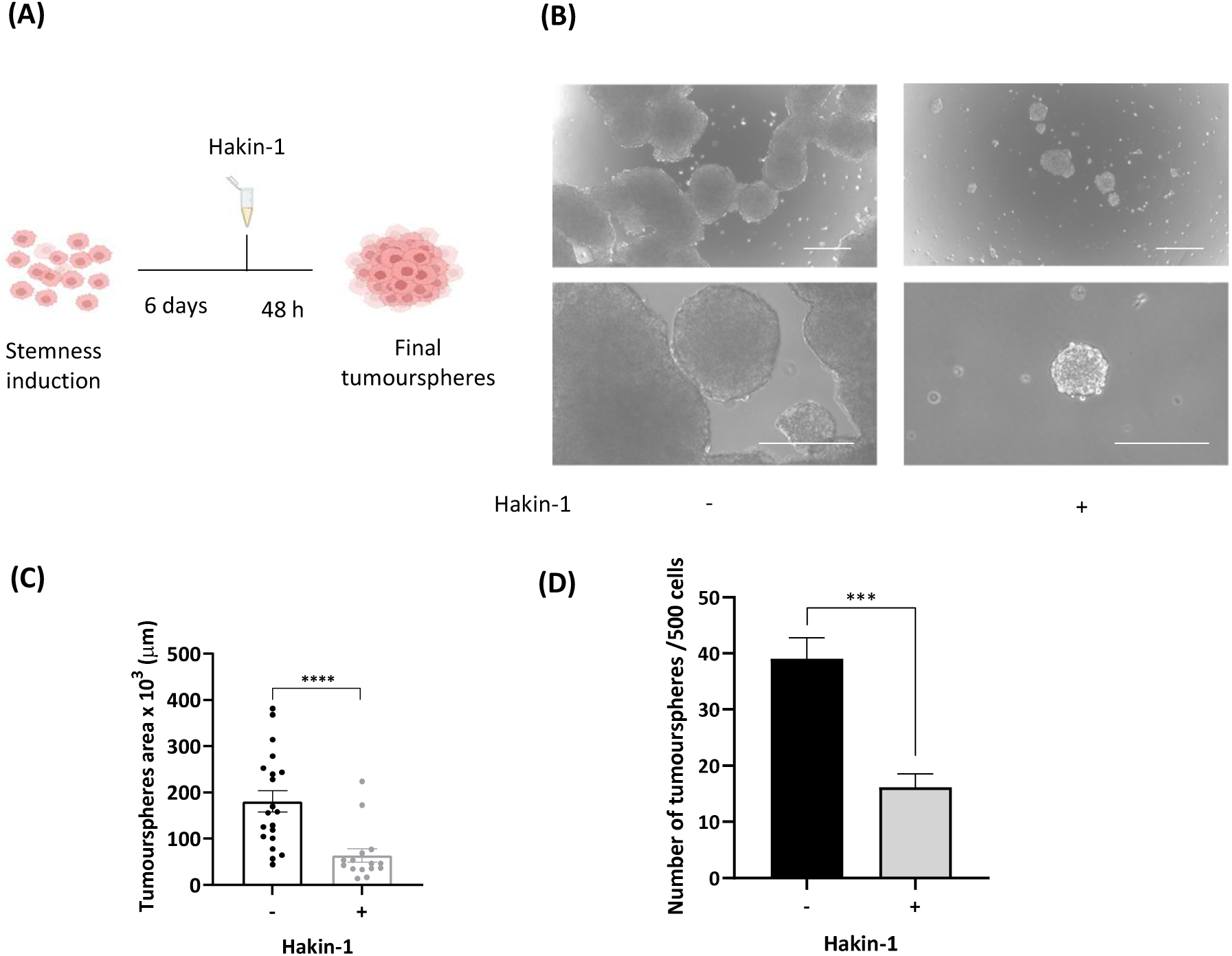
Impact of Hakin-1 treatment on tumourspheres formation. **(A)** Schematic workflow. Tumourspheres already formed were treated with Hakin-1. **(B)** Representative images of tumourspheres in HT29 treated with DMSO (control) or Hakain-1 in day 8 after stemness condition a 4x (upper images, scale bar: 250 μm) or 10x (lower images, scale bar: 200 μm) objective. **(C)** Size of tumourspheres derived from HT29 cells treated with DMSO (control) or 50 μM Hakin-1. Surface area quantification was performed using ImageJ software. Tumoursphere area of at least 15 tumourspheres per experiment were measured. **(D)** Quantification of the number of tumourspheres after DMSO (control) or Hakin-1 treatment by self-renewal assay. Results are expressed as mean ± SEM and statistical analysis was performed using a t-test of GraphPad Prism software (***p < 0.001 and ****p < 0.0001).

**Fig. S4.**
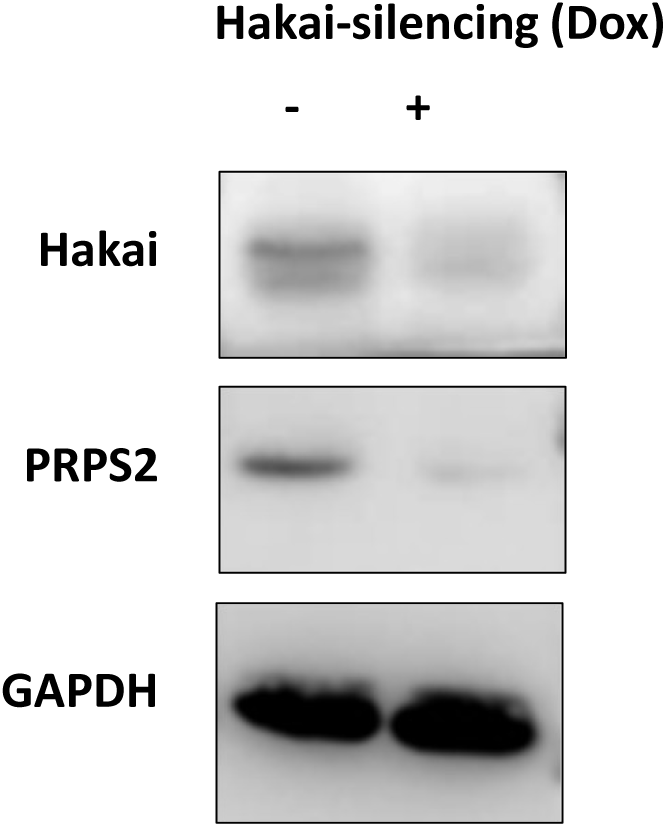
Validation of identified PRPS2 protein as Hakai-regulated protein in tumoursphere by Western Blot. Levels of Hakai and PRPS2 proteins in Hakai-silenced HT29 colon cancer tumourspheres versus control tumoursphere were assessed by Western blot. GAPDH was used as loading control.

## Notes

### Competing Interest Statement

The authors have declared no competing interest.

